# A neural mechanism for discriminating social threat from social safety

**DOI:** 10.1101/2023.07.04.547723

**Authors:** Pegah Kassraian, Shivani K. Bigler, Diana M. Gilly, Neilesh Shrotri, Steven A. Siegelbaum

## Abstract

The ability to distinguish a threatening from non-threatening conspecific based on past experience is critical for adaptive social behaviors. Although recent progress has been made in identifying the neural circuits that contribute to different types of positive and negative social interactions, the neural mechanisms that enable the discrimination of individuals based on past aversive experiences remain unknown. Here, we developed a modified social fear conditioning paradigm that induced in both sexes robust behavioral discrimination of a conspecific associated with a footshock (CS+) from a non-reinforced interaction partner (CS-). Strikingly, chemogenetic or optogenetic silencing of hippocampal CA2 pyramidal neurons, which have been previously implicated in social novelty recognition memory, resulted in generalized avoidance fear behavior towards the equally familiar CS-and CS+. One-photon calcium imaging revealed that the accuracy with which CA2 representations discriminate the CS+ from the CS-animal was enhanced following social fear conditioning and strongly correlated with behavioral discrimination. Moreover the CA2 representations incorporated a generalized or abstract representation of social valence irrespective of conspecific identity and location. Thus, our results demonstrate, for the first time, that the same hippocampal CA2 subregion mediates social memories based on conspecific familiarity and social threat, through the incorporation of a representation of social valence into an initial representation of social identity.

## Main

An animal’s normal functioning in a social community requires that it recognize and discriminate conspecifics based on the valence of its past social encounters with a given individual. The discrimination of past safety- or threat-associated experiences with a conspecific, social fear specificity, is of particular importance for the appropriate choice of approach and avoidance behaviors. Failure to make this distinction may lead to social withdrawal and social anxiety, prominent symptoms of several neuropsychiatric disorders (File and Hyde, 1978; Lissek et al., 2008, 2014; Stein and Stein, 2008; Beckers et al., 2023). Recent studies in rodents have revealed both cortical and subcortical brain regions that assign social valence and regulate appropriate social behaviors (Padilla-Coreano et al., 2022). Most studies have focused on the role of the prefrontal cortex and its subcortical connections, including its importance for formation of social dominance hierarchies (Padilla-Coreano et al., 2022), impairment of social interactions following chronic social defeat (Li et al., 2023) and their enhancement by psychiatric relevant drug treatments (Nardou et al., 2023), and the implementation of conditioned social fear (Xu et al., 2019). However, the neural circuits mediating the memory of past aversive social encounters to enable social fear specificity remain unknown.

One candidate structure for social fear specificity is the hippocampus, whose role in social memory has been well established since the pioneering studies of Brenda Milner and colleagues on patient HM (Scoville and Milner, 1957). More recent studies in rodents have demonstrated, through the exposure of a subject to novel conspecifics, the importance of several subregions of the hippocampus for social novelty recognition memory (SNRM), the ability to distinguish a novel from familiar individual. This includes the hippocampal dorsal CA2 (Hitti, 2014; Stevenson and Caldwell 2014; Smith et al., 2016; Meira et al., 2018; Okuyama, 2016; Meira 2018; Wu et al., 2021), ventral CA1 (Okuyama et al., 2016; Phillips et al., 2019) and ventral CA3 (Chiang et al., 2018) regions, as well as the connections from dorsal CA2 to ventral CA1 (Raam et al., 2017; Meira et al., 2018) and from the supramammilary nucleus to dorsal CA2 (Chen et al., 2020). However, little is known about the neural circuits that enable an individual to distinguish conspecifics based on prior social history, including past threat- or safety-associated experiences. Thus, it is unclear whether the same brain regions that are responsible for the detection of familiarity versus novelty are also important for episodic recollection (Mandler, 1980; Eichenbaum et al. 2007; Yonelinas et al., 2010), e.g. of the valence of past social experiences to generate social fear specificity. Given the known importance of the dorsal CA2 region in SNRM, we here investigate the role of this region in social fear specificity irrespective of social novelty.

As there has been no direct examination of whether mice exhibit social fear specificity, we first developed a modified version of a classic social fear conditioning (SFC) protocol (Toth et al., 2012) to address this issue. In prior SFC studies, application of a footshock to a subject mouse during exploration of a single novel conspecific produced a generalized aversive fear response when the subject was presented one day later with another novel conspecific (Toth et al., 2012; Xu et al., 2019). This is analogous to generalized social fear following social defeat paradigms (Krishnan et al., 2007; Golden et al., 2008; Willmore et al., 2022, Li et al., 2023). To determine whether a subject mouse could show appropriate fear specificity to a threat-associated compared to a safety-associated conspecific, we used a modified version of the SFC paradigm in which a subject mouse simultaneously explored two novel stimulus mice, each confined to wire cup cages, receiving a mild footshock when it explored one stimulus mouse (CS+) but not the other (CS-; Fig. 1a). To probe the memory specificity for the differential conditioning, we re-exposed the subject to the previously encountered CS+ and CS- mice in a novel arena 24 hrs after SFC (Fig. 1a,b). A control group of mice underwent the same protocol and explored the same two stimulus mice as the SFC group but did not receive footshocks during social exploration (Mock-SFC). We used a supervised classification to characterize the behavior of subject mice with a long-short term memory (LSTM) recurrent network with input features constructed based on markerless pose estimation of body parts (Fig. 1a; Methods).

**Fig. 1.**
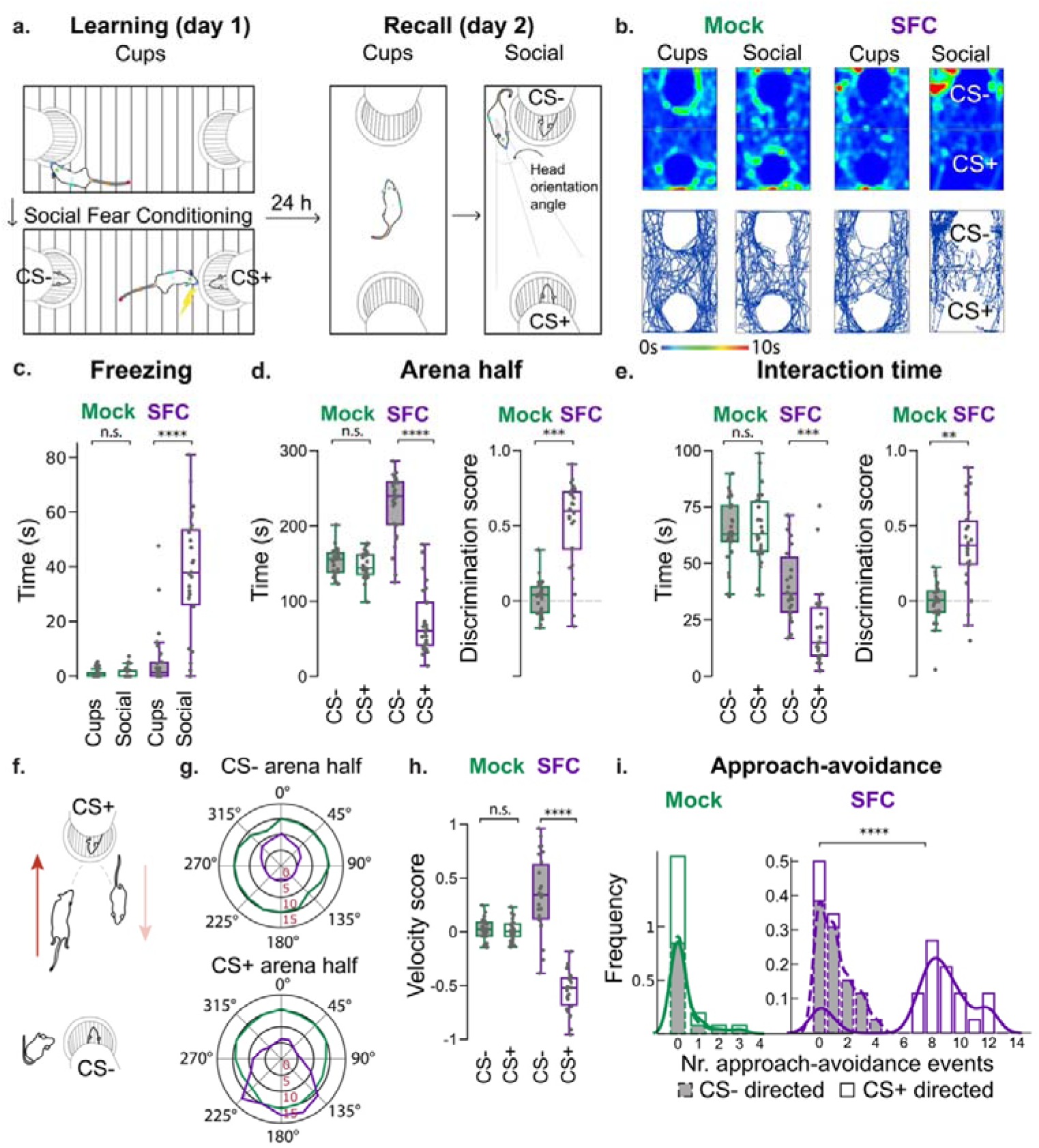
Social fear conditioning (SFC) induces a robust discrimination between safety- and threat-associated conspecifics. **a.** SFC paradigm. Learning (day 1): Empty cup exploration trial followed by SFC trial, with footshock delivered during exploration of CS+ mouse but not CS- mouse. Recall (day 2): Empty cup recall trial (‘Cups’) followed by exposure to CS+ and CS- mouse (‘Social’ recall trial). Novel arena on day 2. Colored dots on subject mouse indicate labels used for pose estimation. Mock-SFC cohort underwent same assay without receiving footshocks. **b.** Example heatmaps and path plots during recall trials for one Mock-SFC and one SFC mouse, illustrating the latter’s preference for the CS- mouse. **c.** Freezing time for Mock-SFC (green) and SFC cohort (purple) during recall trials. SFC cohort displays significantly elevated freezing levels during the ‘Social’ recall trial (two-way repeated measures ANOVA: Cohort x Stage, F(1,100) = 38.82, p<0.0001). **d.** Left plot, SFC cohort spends significantly more time in the arena half containing the CS- than the CS+ mouse (two-way ANOVA: Cohort x Arena half, F(1,100) = 127.44, p<0.0001). Right plot, SFC cohort shows greater discrimination score, (time exploring CS- half minus time exploring CS+ half)/(total exploration time arena), than Mock-SFC cohort (unpaired t-test). **e.** Left, SFC cohort interacted significantly more with the CS- than the CS+ (two-way ANOVA: Cohort x Stimulus mouse, F(1,100) = 10.91, p<0.01). Right, discrimination score based on interaction time (unpaired t-test). **f.** Illustration of approach-avoidance behaviors (red arrow: approach, pink arrow: avoidance). **g.** Speed per head orientation angle towards the CS- (left) and the CS+ mouse (right) within the respective arena half. The same analysis is performed for the Mock-SFC cohort for a randomly selected stimulus mouse. Approach-avoidance behaviors are classified by a LSTM neural network. **h.** Relative velocity of approach compared to retreat towards CS- or CS+ mouse. Velocity score = [(approach velocity – retreat velocity)]/[sum of velocities], two-way ANOVA: Cohort x Arena half, F(1,100) = 117.05, p<0.0001. **i.** Frequency histograms showing greater number of approach-avoidance behaviors towards CS+ than CS- mouse for the SFC but not the Mock-SFC cohort. Kolmogorov-Smirnov test: **** p<0.0001. N = 13 females plus 13 males per cohort. Box plots: central line, median; bottom and top edges, 25th and 75th percentiles; whiskers, most extreme data points (excluding outliers); dots, individual animals. Bonferroni post-hoc and t-tests: ** p<0.01, *** p<0.001, **** p<0.0001, n.s.: not significant. No significant effect of sex was observed.

We found that the SFC protocol produced in both female and male mice robust social fear behaviors when tested 24 hours after conditioning. Thus, the subject mouse displayed significantly greater freezing behavior during the recall session compared to the Mock-SFC group (Fig. 1c). As previously reported for the standard SFC protocol, the SFC-induced fear response was highly selective for the social context (Toth et al., 2012; Xu et al., 2019), as neither group showed freezing behavior during exposure to the empty cups in the novel context. Moreover, the groups did not differ in general anxiety measured by the elevated plus maze or in locomotor activity (Extended Data Fig. 1a,b). Importantly, we found a high degree of specificity of social fear behavior with our SFC paradigm, in which the SFC mice clearly discriminated the CS+ from the CS- stimulus mice (Fig. 1b,d-i). Thus, SFC subject mice interacted to a significantly greater extent with the CS- compared to the CS+ and spent a greater fraction of time in the half of the arena containing the CS- (Fig. 1b,d,e). Moreover, the SFC mice showed a much greater incidence of approach-avoidance behaviors (Xu et al., 2019) to the CS+ than the CS- (Fig. 1 f-i). The Mock-SFC cohort in contrast did not display differential responses toward the two conspecifics and rarely displayed approach-avoidance behaviors.

To determine whether the CS- mice were viewed as a neutral stimulus or had a positive, safety-associated valence, we examined SFC using a three-chamber recall arena, in which the two stimulus mice were confined to wire cup cages in the end chambers of the arena and the middle chamber was empty. We found in independent cohorts that social fear conditioned mice spent most of their time in the CS- chamber as opposed to either the CS+ or the ‘neutral’ central chamber, suggesting that the non-reinforced interaction partner might signal safety (Extended Data Fig. 2a,b). These results thus provide to our knowledge one of the first demonstrations that mice can discriminate a threat-associated from a safety-associated conspecific. Moreover, the display of approach-avoidance during recall in a novel arena and, therefore, independent of spatial or contextual cues, suggests the distinct association of the aversive stimulus with conspecific identity.

**Fig. 2.**
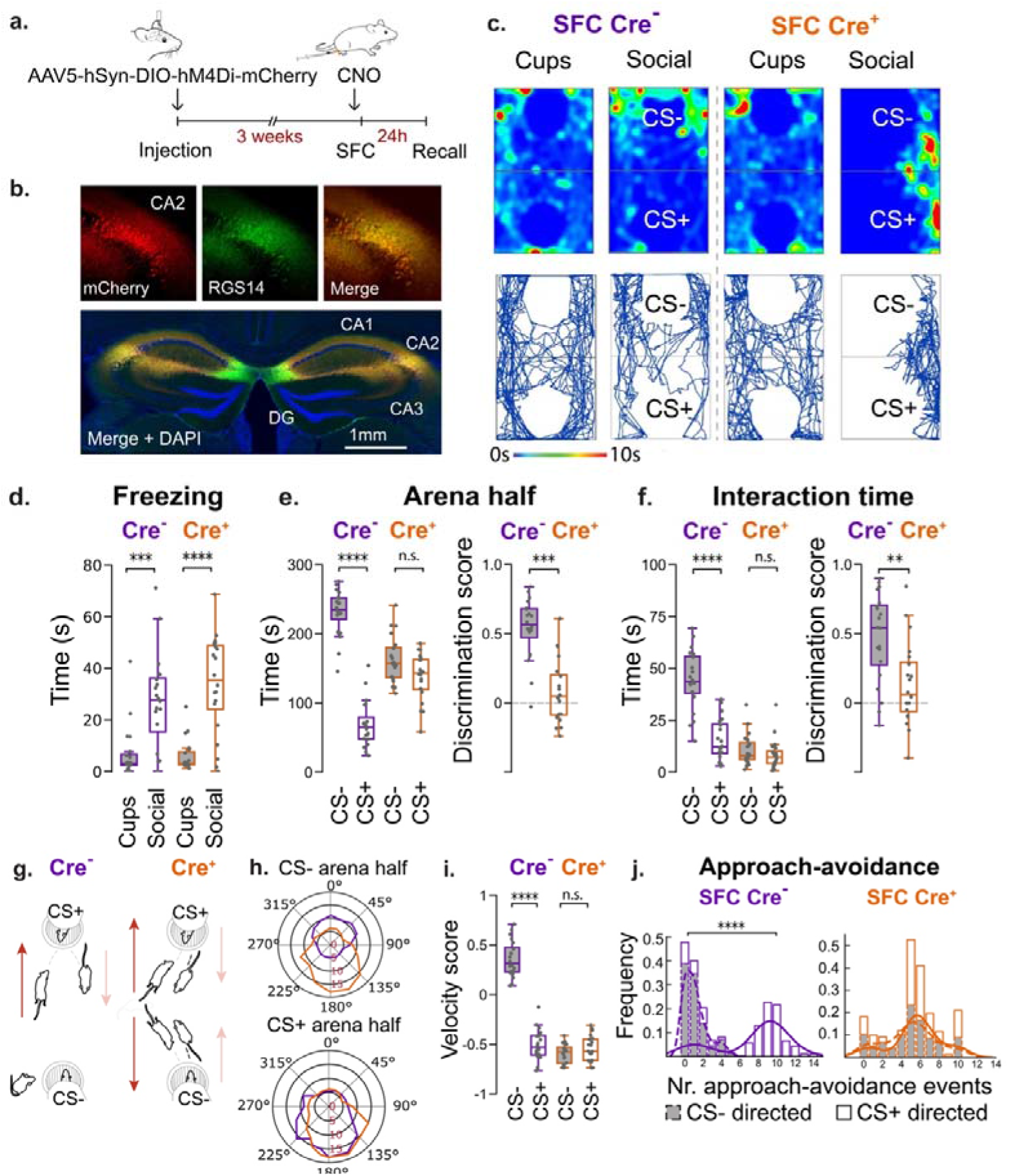
CA2 silencing during SFC learning caused a loss of social fear specificity. **a.** AVPR1b-Cre+ and Amigo2-Cre+ and wild-type littermate cohorts are injected with AAV2/5-hSyn.DIO.hM4D(Gi)-mCherry in CA2. After 3-4 weeks, cohorts are administered intraperitoneally CNO 30 min before SFC on day 1. **b.** Confocal images of an example coronal section from an Amigo2-Cre mouse showing selective expression of mCherry-tagged hM4Di in CA2 pyramidal cells, identified by co-expression of the CA2 marker protein RGS14. **c.** Example heatmaps and path plots for one SFC Cre- and one SFC Cre+ mouse during recall trials (no CNO), showing selective avoidance of the CS+ mouse by Cre- mouse and the non-selective avoidance of both CS+ and CS- mice by Cre+ mouse. **d.** Freezing time for Cre- (purple) and Cre+ cohort (orange) during ‘Cups’ and ‘Social’ recall trials. Both cohorts display similar freezing levels when comparing ‘Cups’ recall trials and when comparing ‘Social’ recall trials (two- way repeated measures ANOVA: Cohort x Stage, F(1,76) = 1.3846, p>0.05) and significantly greater time freezing during the ‘Social’ than the ‘Cups’ recall trial (Bonferroni post-hoc tests). **e.** Left, SFC leads to increased time in arena half containing CS-mouse for Cre- cohort; Cre+ cohort spends equivalent time in arena halves (two-way ANOVA: Cohort x Arena half, F(1,76) = 82.93, p<0.0001). Right, discrimination score based on time in CS- versus CS+ arena half significantly greater in Cre- than Cre+ (unpaired t-test). **f.** Left, Cre- but not Cre+ cohort shows greater interaction time with CS- than CS+ mouse (two-way ANOVA: Cohort x Stimulus mouse, F(1,76) = 32.92, p<0.0001). Right, discrimination score based on interaction time (unpaired t-test). **g.** Cre- mice exhibit slow approach to the CS+ followed by rapid retreat to the CS- half (left). Cre+ mice display approach- avoidance towards both the CS+ and CS- (right). **h.** Polar plot for speed per head orientation angle towards the CS- (left) and the CS+ (right) within the respective arena half, illustrating the generalized avoidance behaviors of the SFC Cre+ cohort. **i.** Velocity score shows preferential approach of CS- compared to CS+ for Cre- but not Cre+ cohort (two-way ANOVA: Cohort x Arena half, F(1,76) = 200, p<0.0001). **j.** Frequency histogram for approach avoidance behaviors show significantly greater number of approach-avoidance behaviors towards the CS+ than towards the CS- by Cre- but not Cre+ mice. Kolmogorov-Smirnov test: **** p<0.0001. N = 10 females plus 10 males per cohort. Avpr1b-Cre+ and Amigo2-Cre+ numbers balanced across cohort and sex with no significant effect of Avpr1b-Cre+ versus Amigo2-Cre+ genotype and sex observed. Bonferroni post-hoc and t-tests: ** p<0.01, *** p<0.001, **** p<0.0001, n.s.: not significant.

### The CA2 region is necessary for the dissociation of a safety- from a threat-associated conspecific

The observation that mice of both sexes can robustly discriminate between a safety- and a threat-associated conspecific allowed us to probe if the same brain region required for social novelty recognition memory also plays an important role for this discrimination. Because of the known role of CA2 PNs in social novelty recognition memory, we addressed this question using chemogenetics to silence CA2 PNs during SFC. We selectively expressed the inhibitory G-protein coupled hM4Di receptor (iDREADD) in CA2 PNs by injecting Cre-dependent AAV2/5 into dorsal CA2 of female and male Amigo2-Cre+ (Hitti and Siegelbaum, 2014) or Avpr1b-Cre+ (Williams Avram et al., 2019) mice. Both mouse lines express Cre recombinase relatively selectively in CA2 and enable CA2- specific expression using targeted Cre-dependent AAV injections into dorsal CA2. We used the two mouse lines, the Amigo2-Cre being a transgene and the Avpr1b-Cre a knock-in, to strengthen any conclusions that any deficits in behavior were due to specific effects on CA2 PN activity, rather than non-specific or off-target actions. Approximately 3-4 weeks after viral injection, iDREADD-expressing Cre+ mice and a balanced number of Cre- littermate controls were intraperitoneally injected with the iDREADD agonist clozapine-N-oxide (CNO, 5 mg/kg) 30 minutes prior to SFC and then tested for social fear recall 24 hours later in the absence of CNO (Fig. 2a-c). In a separate set of control experiments, we compared uninjected Cre+ mice with Cre- mice in the presence of CNO to confirm than any change in behavior was not due to genotype.

We found that both CA2-silenced Cre+ mice and Cre- mice exhibited a marked social fear during the recall session after SFC, as assessed by their increased freezing behavior in the presence of both CS+ and CS- (Fig. 2d). Strikingly, the CA2-silenced Cre+ mice no longer discriminated between the CS- and the CS+, spending a similar amount of time in the CS- and CS+ halves of the arena and a similar time actively exploring both stimulus mice (Fig. 2c,e,f). In contrast the Cre- control group displayed a normal preference for the CS- as opposed to the CS+. Moreover, whereas the Cre- control group showed selective approach-avoidance behavior to the CS+, the Cre+ group showed a generalized fear response to both the CS+ and CS-, with each eliciting similar elevated levels of approach-avoidance behavior (Fig. 2g-j). These results in an open arena were confirmed in a three-chamber recall arena, where Cre+ mice spent most of the recall session in the middle chamber, avoiding both the CS- and the CS+ chamber. In contrast, Cre- mice showed a normal preference for the CS- chamber (Extended Data Fig. 2c-e). In the absence of iDREADD expression, the Cre+ and Cre- mice showed similar SFC responses with a high level of specificity for the CS+ relative to the CS-, demonstrating that the difference in behaviors in the iDREADD-injected Cre+ and Cre- groups was due to iDREADD-mediated CA2 silencing, rather than differences based on genotype (Extended Data Fig. 3).

**Fig. 3.**
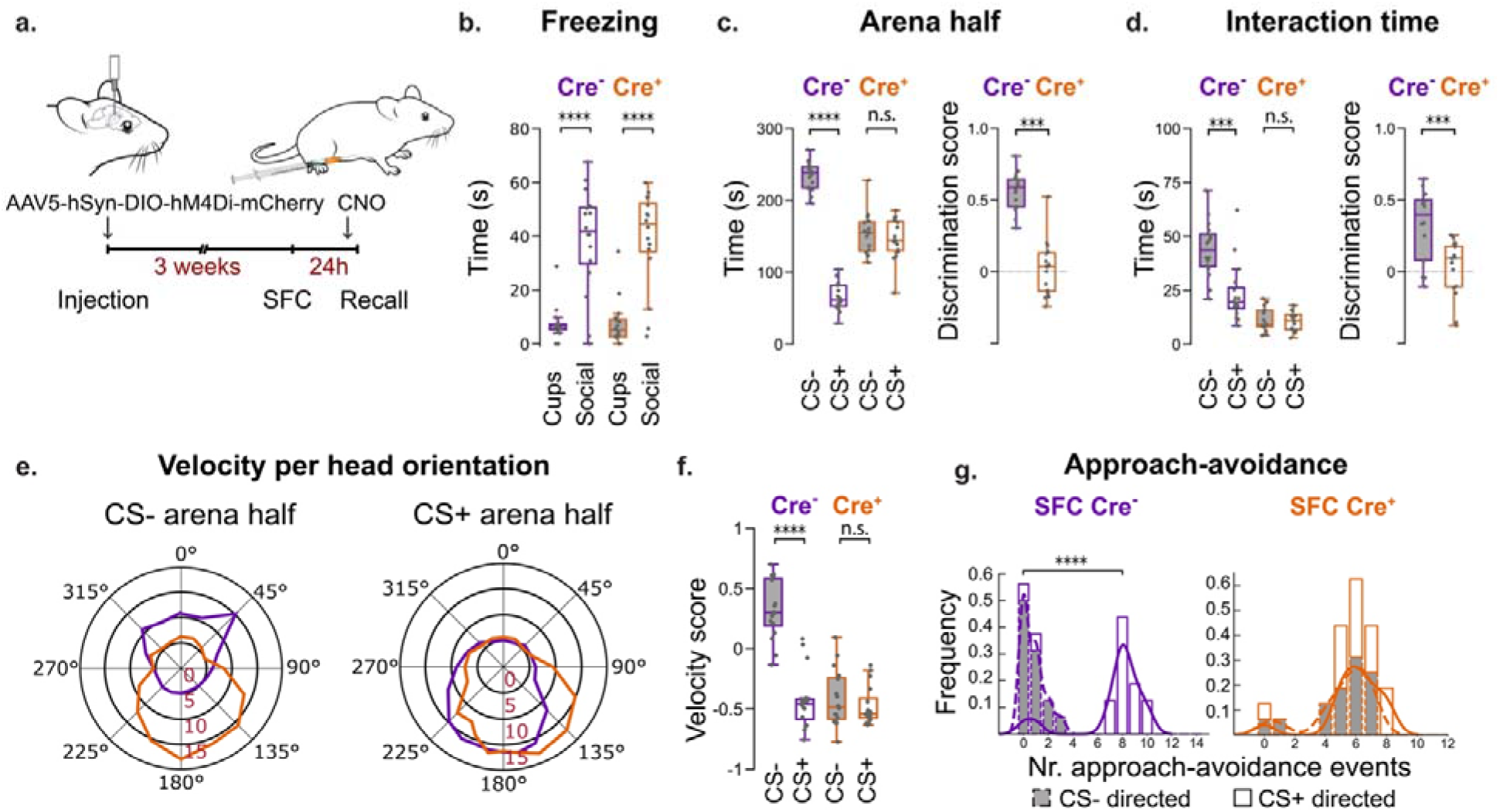
CA2 silencing during social fear recall also leads to a loss of social fear specificity. **a.** Amigo2-Cre+ and AVPR1b- Cre+ mice and their Cre- littermates are injected in CA2 with AAV2/5-hSyn.DIO.hM4D(Gi)-mCherry. After approximately 3-4 weeks, both cohorts undergo SFC in absence of CNO during day 1 Learning session. 30 min prior to recall on day 2, both groups are administered CNO intraperitoneally. **b**. Freezing during recall after SFC. Cre+ and Cre- cohorts display similar freezing levels when comparing ‘Cups’ recall trials and ‘Social’ recall trials (two-way repeated measures ANOVA: Cohort x Stage, F(1,60) < 0.0001, p>0.05) and significantly greater time freezing during the ‘Social’ than the ‘Cups’ recall trial (Bonferroni post-hoc tests). **c.** Cre- mice but not Cre+ mice selectively avoid the CS+ arena half, measured either by exploration time (left, two-way ANOVA: Cohort x Arena half, F(1,60) = 168.02, p <0.0001) or discrimination score (right, unpaired t-test). **d.** Cre- mice preferentially interact with CS- relative to CS+ whereas Cre+ mice interact similarly with both, measured by interaction time (left, two-way ANOVA: Cohort x Stimulus mouse, F(1,60) = 15.46, p <0.0001) or discrimination score (right, unpaired t-test). **e.** While SFC Cre- mice display slow approach towards the CS+ followed by rapid retreat to the CS- half (left), SFC Cre+ mice display approach-avoidance behaviors towards both the CS+ and CS- (right). **f.** Velocity score. Cre- but not Cre+ mice show higher velocity of approach to Cs- than CS+, two-way ANOVA: Cohort x Arena half, F(1,60) = 37.45, p<0.0001. **g.** Frequency histogram shows Cre- group has significantly greater number of approach-avoidance behaviors targeted at the CS+ than CS-; Cre+ cohort displays a similar number of approach-avoidance behaviors towards the CS+ and the CS- (right). Kolmogorov-Smirnov test: **** p<0.0001. N = 8 females plus 8 males per cohort. Avpr1b-Cre+ and Amigo2-Cre+ numbers balanced across cohort and sex with no significant effect of Avpr1b-Cre+ versus Amigo2-Cre+ genotype and sex observed. Bonferroni post-hoc tests: ***p < 0.001, ****p < 0.0001, n.s.: not significant.

Importantly, the effect of CA2 silencing to cause an indiscriminate social fear response during recall is not due to a general increase in anxiety or non-specific fear responses as Cre+ and Cre- cohorts spent equal times: 1. Freezing during the non-social ‘Cups’ stage of recall, 2. Exploring the open arms in the Elevated Plus Maze, and 3. Engaging in locomotor activity (Figs. 2d, 3b; Extended Data Figs. 1c,d). Moreover, the CA2-silenced mice did not show freezing behavior in a recall session with empty cups. Thus, the non-selective fear response is highly specific to a social context.

As CNO injection can inhibit the activity of neurons expressing iDREADD for several hours, we next used an optogenetic approach to silence CA2 selectively during the five minutes of the SFC trial. We used Cre-dependent AAV injection to express Arch3.0 in CA2 of Amigo2-Cre and Avpr1b-Cre mice and their Cre- littermates. We used a fiber optic probe impanted over CA2 to illuminate this region with yellow light continuously during the SFC trial. Similar to the chemogenetic results, we found that optogenetic silencing of CA2 during social fear encoding disrupted the discrimination between safe and threat-associated conspecifics during social fear recall in Cre+ but not Cre- mice (Extended Data Fig. 4).

**Fig. 4.**
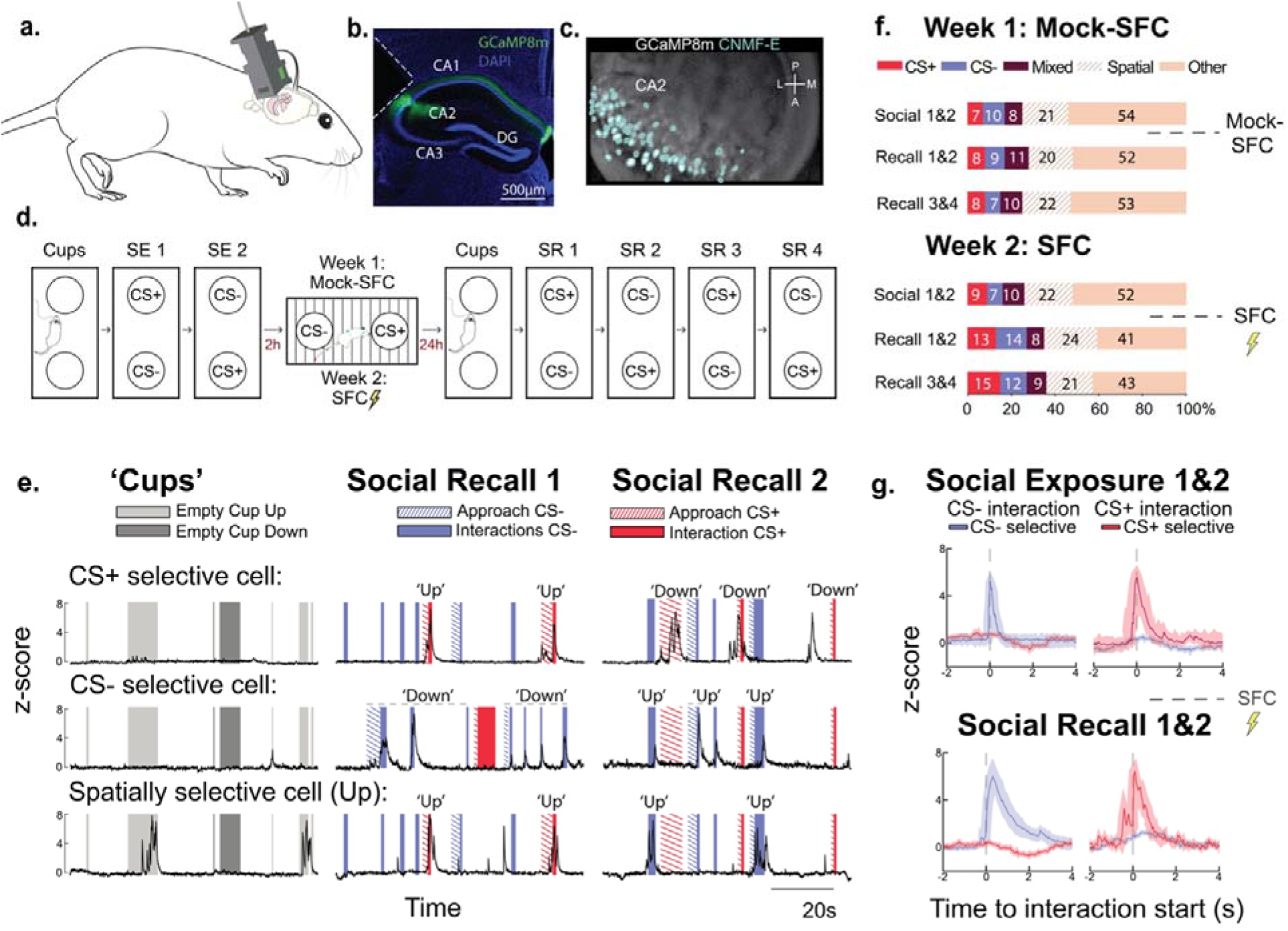
Calcium imaging of CA2 pyramidal cells before and after SFC. **a.** CA2 calcium activity is imaged in freely behaving mice during an extended version of the SFC paradigm (panel d) using a microendoscope with the Cre-dependent Ca^2+^ indicator GCaMP8m expressed in CA2 pyramidal cells of Amigo2-Cre or Avpr1b-Cre mice. **b.** Confocal image of GCaMP8m expression in CA2 pyramidal cells. Dashed lines indicate the GRIN lens tract. **c.** Maximum intensity projection of the field of view and cell detection with CNMF-E from an example imaging session. **d.** SFC paradigm. On day 1, a subject mouse was first allowed to explore an open arena (the same used above for SFC recall) containing two empty cups for 5 min (‘Cups’), followed by two 5- min trials in which the subject explored the prospective CS+ and CS- mice (SE, ‘Social Exploration’ trials), with positions swapped in each trial. In week 1, SE trials were followed 2 hours later by a Mock-SFC trial in the SFC chamber. On day 2, the subject was re-exposed to the open arena with empty cups (‘Cups’), followed by four consecutive ‘Social Recall’ (SR) trials, with positions of stimulus mice swapped between each trial. In week 2, the same procedure was repeated with two novel stimulus mice and with the Mock-SFC trial replaced by a SFC session. **e.** Representative GCaMP8m z-score activity traces after SFC of a CS+ selective cell (top row), a CS- selective cell (middle row) and a spatially selective cell (bottom row) during interactions with empty cups (left column) or a stimulus mouse during the two social recall trials (middle and right columns). ‘Up’ and ‘Down’ legends: social interactions with a stimulus mouse in the upper or lower cup. Note positions of stimulus mice swapped between SR trials. Cells whose activity during social exploration with the same stimulus mouse differed significantly from baseline activity (pL≤L0.01) during two consecutive social recall trials and across both spatial positions (upper and lower cup) were classified as CS+ and/or CS- selective, according to the nature of the responses. Cells which were both CS+ and CS- selective were classified as ‘mixed selective’. Cells whose activity during social explorations was significantly different from baseline across one spatial position independent of stimulus mouse identity were classified as spatially selective, and else cells were classified as ‘Other’. **f.** Fraction of CA2 cells with indicated response profiles; N = 10 mice. SFC significantly increased the fraction of cells that responded to the CS+ or the CS- (Chi-square test; χ (6) = 34.382, **** p<0.0001); no change of the CS-selective population was observed for the Mock-SFC cohort (χ ^2^(6) = 4.943, p = 0.551). **g.** Average z-scored responses for CS- and CS+ selective cell populations before and after SFC (mean±SEM across N=5 Amigo2-Cre mice plus N = 5 Avpr1b-Cre mice) aligned to the start of CS- (left) or CS+ (right) interactions (dashed line).

The above results demonstrate the importance of CA2 for the encoding of social fear memory. To investigate if CA2 activity during recall of a social fear memory is required for the discrimination between safe and threat-associated conspecifics, independent Cre+ and Cre- cohorts injected with Cre- dependent iDREADD AAV underwent SFC in the absence of CNO. We then injected the mice with CNO 30 min prior to social fear recall 24 hrs after SFC (Fig. 3a). Similar to when we silenced CA2 during learning, both Cre+ and Cre- mice displayed increased freezing during social recall (Fig. 3b). However, CA2-silencing during recall suppressed the preference of the subject to explore the CS- relative to the CS+ (Fig. 3c,d), with the subject displaying a similar amount of approach-avoidance behaviors directed towards both CS+ and CS- stimulus mice (Fig. 3e-g). In contrast, the Cre- cohort maintained its preference for the CS- conspecific.

In principle, the subject mice might avoid the CS+ during SFC recall if the CS+ emitted an alarm signal in response to having witnessed the subject’s distress during SFC. As the Mock-SFC group interacted with the same CS+ and CS- during the recall trial as did the SFC group, any alarm signal must have been selectively emitted by the CS+ in response to the SFC mice. However, this scenario is unlikely since when we expressed iDREADD and silenced CA2 in the two stimulus mice with CNO injection 30 minutes prior to SFC we found neither an effect on SNRM nor on the ability of the SFC mice to discriminate between the CS+ and CS- stimulus mice (Extended Data Figs. 5,6).

**Fig. 5.**
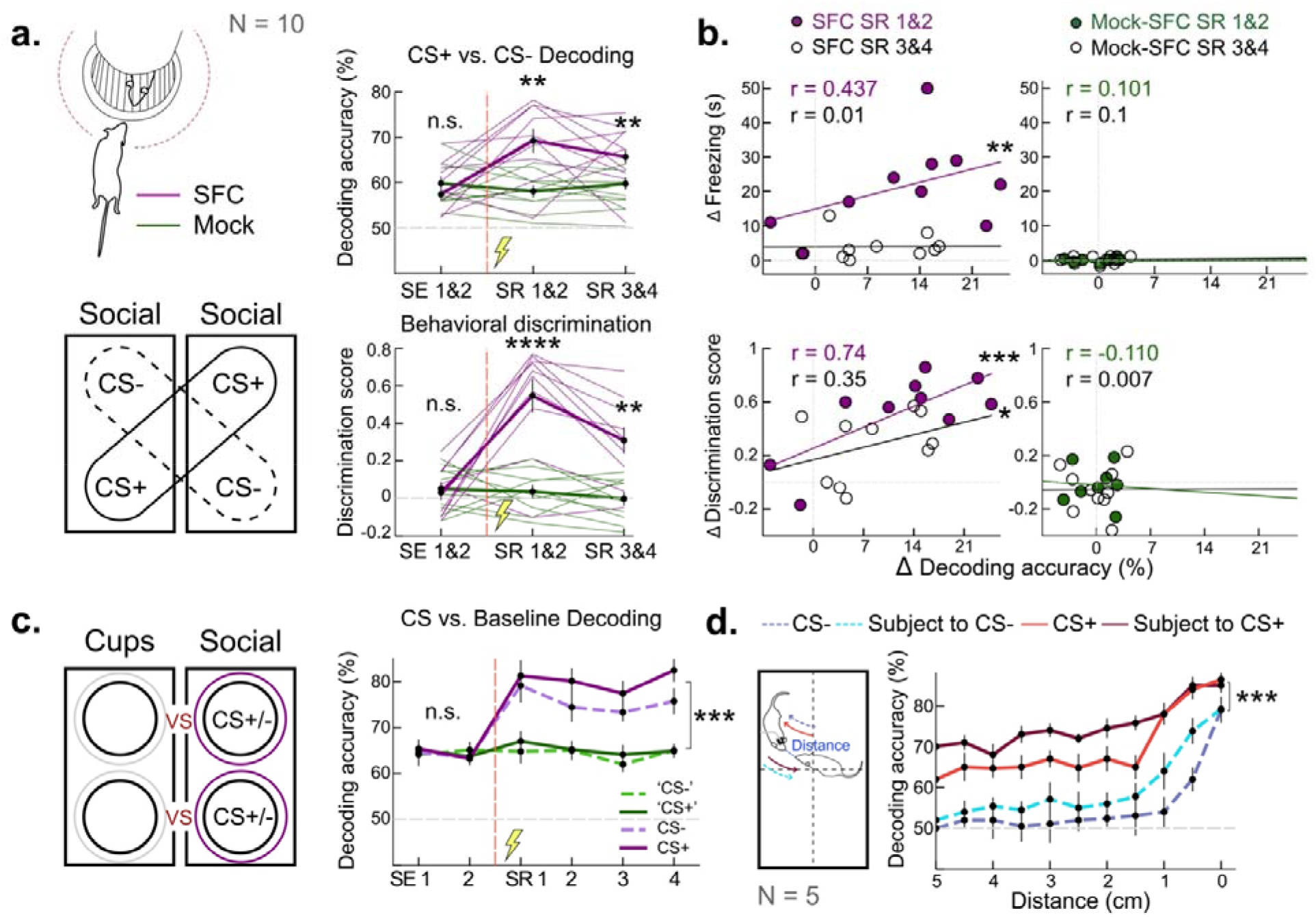
CA2 encodes representations of a safe versus a threat-associated conspecific. **a.** Left column: Calcium data obtained during social interactions is pooled across two spatial positions in the two SR trials to balance spatial information and train a linear classifier to discriminate CS+ versus CS- interactions. Right column: Decoding accuracies (top) and behavioral discrimination (bottom) scores (both averaged across two consecutive trials) are significantly greater for the SFC (purple) as compared to the Mock-SFC (green) cohort during SR trials after SFC. Two-way ANOVA: Cohort x Session, F(2,54) = 8.4, p<0.001 for decoding and F(2,54) = 11.58, p<0.0001 for behavior. Bonferroni post-hoc tests: ** p<0.01, **** p<0.0001. **b.** Changes in decoding accuracy and behavioral discrimination before and after SFC are highly correlated (left panel); no such correlation is observed after Mock-SFC (right panel). Spearman’s rank correlation coefficient: * p<0.05, ** p<0.01, *** p<0.001. **c.** Decoding accuracies of CS+ (solid) or CS- (dashed) versus empty cup in same arena position are significantly enhanced after SFC (purple) but not after Mock-SFC (green). Two-way ANOVA: Cohort x Session, F(5,108) = 4.717, p<0.01. Bonferroni post- hoc tests: **** p<0.0001 for SR trials of Mock-SFC vs. SFC cohorts. **d.** Open arena assay where the subject mouse is exposed to freely behaving CS+ or CS- in consecutive recall sessions 24 hrs after SFC. Approach of subject by CS+ (red) or of subject to CS+ (purple) can be decoded for a greater range of distances and with higher accuracies than for approach of subject by CS- (dashed blue) or of subject to CS+ (dashed cyan) throughout all considered distances (two-way ANOVA: Approach type x Distance, F(30,176) = 2.863, p<0.01). Bonferroni post-hoc tests: *** p<0.001 for CS+ vs. CS- decoding accuracies and all distances considered. N.s.: not significant. Bold lines and bars in panels a, c, d: mean±SEM across cohorts. N = 10 mice with a total of 514 cells for panels a, b, c, and an independent cohort of N = 5 Amigo2-Cre with 382 cells mice for panel d.

**Fig. 6.**
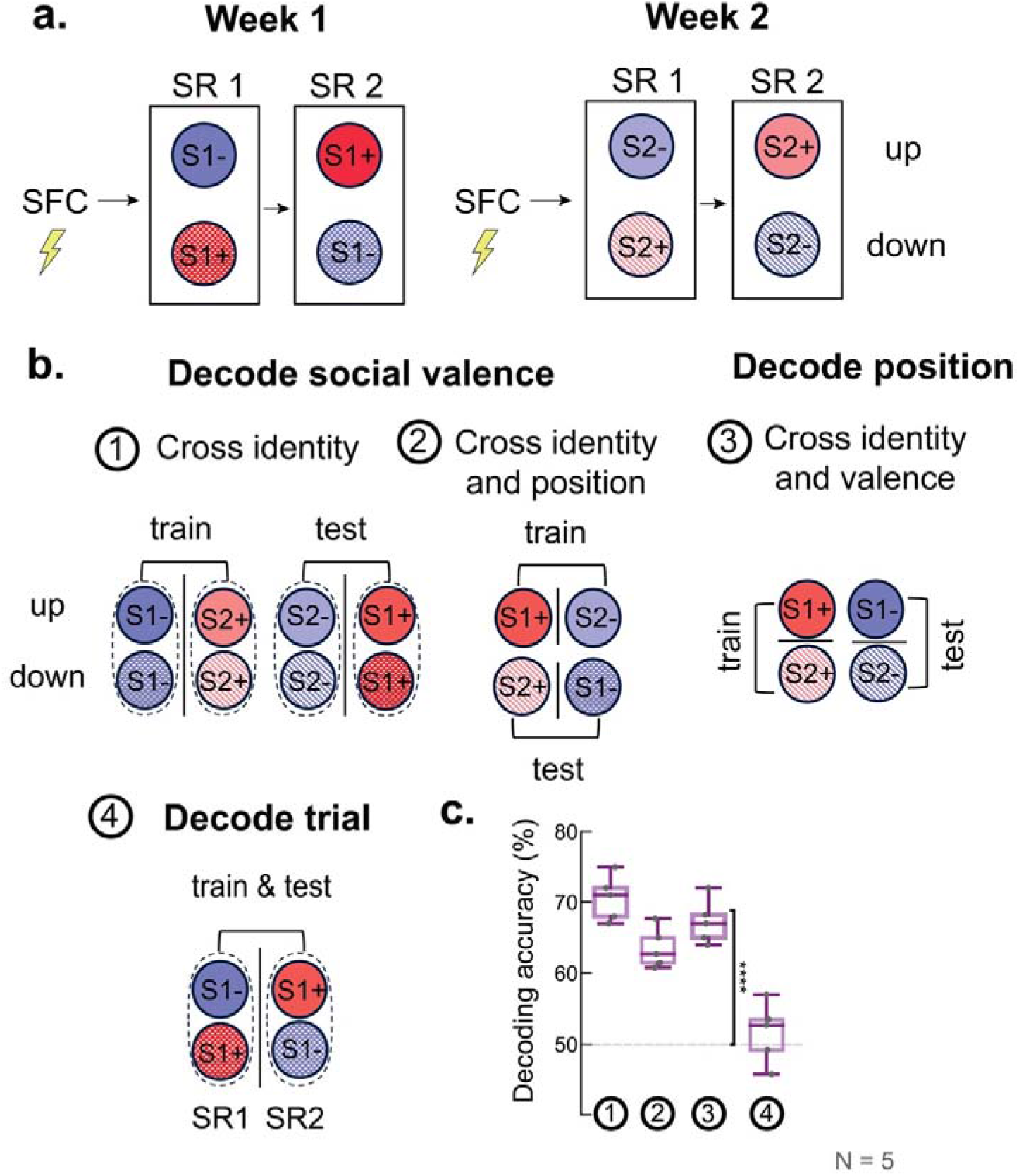
The CA2 region encodes a generalized or abstract representation of social valence. **a.** Experimental timeline: Two SFC sessions, separated by one week, with stimulus mice CS_1_+ and CS_1_- on week 1 and CS_2_+ and CS_2_- on week 2. CS+ mice colored blue and CS- mice colored red. Up position, solid color; Down position, cross-hatched. Week 1 mice, darker shades; Week 2 mice, lighter shades. **b.** (1) Cross-identity social valence CCGP is calculated by a classifier trained to dissociate one pair of CS+ and CS- mice (eg, CS1- versus CS2+) and tested on the second pair of stimulus mice not used for training (eg, CS2- versus CS1+). Training and testing pairs are then swapped and decoding accuracies averaged. For each decoding test, data from up and down positions are combined. (2) Cross-identity and cross-position social valence CCGP is calculated with the additional constraint that training and testing data are obtained from social interactions with stimulus mice in distinct spatial positions (i.e., upper vs. lower cup). Decoding accuracy averaged across the four possible train-test schemes. (3) Cross-identity position CCGP, where the decoder is trained to dissociate between social interactions around the upper versus lower cup, and tested on the upper versus lower cup with stimulus mice of the opposite valence. Decoding accuracy averaged across the four possible train-test schemes. (4) Bottom right: Binary decoding of SR1 versus SR2 recall trials, resulting in a total of two possible decoding schemes (week 1 and week 2). **c.** CCGP decoding results averaged from individual mice: Cross-identity Social Valence CCGP = 70.6 ± 1.4% (mean ± SEM throughout figure); Cross-identity and Cross-position Social Valence CCGP = 63.5 ± 1.3%; Position CCGP = 67.2 ± 1.3%; Trial decoding = 51.6 ± 1.9%. Paired t-tests against shuffled data: *** p<0.001; ns, p > 0.05. N = 5 Amigo2-Cre mice with a total of 245 cells tracked across both SFC sessions. See Methods for a detailed discussion of the CCGP decoding analysis.

Previous studies on novelty recognition memory found that CA2 was selectively required for social novelty recognition and not for novel object recognition memory tasks (Hitti and Siegelbaum, 2014; Oliva et al., 2020). To assess whether the role of CA2 in the dissociation of safety- and threat-associated stimuli is specific to social fear, we implemented an analogous object fear conditioning protocol (OFC) in which mice received a foot shock only when interacting with one of two distinct novel objects (Extended Data Fig. 7). Mice successfully acquired avoidance of the foot-shock-paired object. Chemogenetic silencing of CA2 during the OFC learning trial did not impair the discrimination between the CS+ and CS- objects during the recall trial. This finding is in line with previous studies showing that CA2 silencing does not impair object recognition memory (Hitti and Siegelbaum, 2014; Oliva et al., 2020) or the formation of reward-associations with non-social odors (Hassan, Bigler and Siegelbaum, 2023).

In summary, silencing CA2 pyramidal neurons with chemo- or optogenetic methods prevented the discrimination of threat-associated from safety-associated individuals, resulting in generalized avoidance fear behavior towards both CS- and CS+ mice. Thus, social fear discrimination, which requires CA2, is dissociable from the association of threat to a social context, which is independent of CA2. Moreover, the selectivity of CA2 for social fear is further supported by our finding that this region is not required for the discrimination of safety- and threat-associated inanimate objects.

### CA2 pyramidal cells selectively respond to distinct conspecifics

Motivated by our finding that CA2 activity was required for the dissociation of safety- and threat-associated conspecifics, we investigated whether CA2 neural activity differentially encoded the CS+ and CS- mice. To this end, we expressed the calcium indication GCaMP8m in CA2 pyramidal neurons and used a head-mounted miniature microendoscope to record calcium dynamics of individual CA2 neurons in freely behaving mice before and after SFC (Fig. 4a-d). On day 1, prior to SFC, a subject mouse was first allowed to explore an open arena (the same used above for SFC recall) containing two empty cup cages for 5 minutes. This was immediately followed by two 5-minute trials in which the subject explored the prospective CS+ and CS- mice, with their locations swapped in each trial. These ‘Social Exploration’ trials were followed 2 hours later by a Mock-SFC trial in the SFC chamber. On day 2, we re-exposed the subject to the open arena with empty cups, followed by four consecutive ‘Social Recall’ trials, with the positions of the stimulus mice swapped between each ‘Social Recall’ trial. As CA2 encodes both social and spatial information (Mankin et al., 2015; Alexander et al., 2016; Donegan et al., 2020; Oliva et al., 2020; Boyle et al., 2022), swapping the position of the mice allowed us to dissect CA2 responses to the social and spatial variables separately. One week later, the same procedure was repeated with two novel stimulus mice, except we subjected mice to the SFC protocol as described above. This extended Mock-SFC/SFC protocol permitted us to assess changes in CA2 representations of conspecifics in the same subject mouse following a neutral compared to a threat-associated experience with a conspecific.

We first assessed the effect of SFC on the response profile of individual CA2 pyramidal cells across 10 mice, with an average of 65.2L±L10 simultaneously imaged cells per mouse (mean ± SD). When we assessed CA2 activity prior to the Mock-SFC during a neutral Social Exposure trial, we found that 7.0 ± 2.8% (mean ± SEM) and 10.0 ± 3.3% of the active CA2 neurons responded selectively to the prospective CS+ and CS-, respectively, with 8.0 ± 3.8% of CA2 responding to both mice (see Methods). There was no significant change in these proportions of responsive cells 24 hours after the Mock-SFC trial (Fig. 4e,f), with 8 ± 2.8% of cells responding selectively to the CS+, 9 ± 3.4% to the CS-, and 11 ± 3.7% responding to both. In marked contrast, the fraction of cells that responded selectively to the CS+ and the CS- increased significantly after SFC, to 13 ± 4.1% and 14 ± 3.7% of cells, respectively, representing a 64% increase in the fraction of cells selective for the CS+ and an 86% increase in cells selective for the CS- after SFC relative to the fraction of selective cells before SFC (Population of CS+ or CS- selective cells; Chi-square test; χ^2^(6) = 34.382, **** p<0.0001). There was no change in the fraction of cells responding to both mice (χ^2^(6) = 4.943, p = 0.551). When we aligned the responses of the CS+ and CS- cells to the start of each social interaction bout, the event-triggered average for each group had a peak activity near the start of the interaction (Fig. 4g). The CS+ selective cells displayed an increase in Ca^2+^ responses slightly prior to the start of an interaction bout, during the approach of the subject to the CS+ stimulus mouse.

### CA2 population activity encodes a memory of a safety- and threat-associated conspecific

Previous work has shown that CA2 population activity encodes both the novelty and the identity of conspecifics (Donegan et al., 2020; Oliva et al., 2020; Boyle, Posani et al., 2022). Although the role of the hippocampus for general associative learning is well-established (Basu and Siegelbaum, 2015; Moser et al., 2017; Buszáki et al., 2022) and the hippocampus has been found to incorporate threat-associated experiences in non-social odor representations in head-fixed mice (Biane et al., 2023), there have been no studies to date examining whether hippocampus contains a representation of past threat-associated experience with conspecifics of similar degrees of familiarity. Moreover, we are aware of no studies that address whether any form of socially-related experience is incorporated in hippocampal representations in freely moving mice.

To address such questions, we used a linear Support Vector Machine (SVM) to assess the extent to which CA2 activity can discriminate the CS+ from the CS- before and after SFC. To construct binary classes for the CS+ and CS- independent of spatial location, we pooled calcium responses obtained during interactions with a given stimulus mouse around both the upper and the lower cup from two consecutive trials (Fig. 5a, upper left). In line with previous work (Boyle, Posani et al., 2022), we found that conspecific identity was decoded from CA2 activity at levels significantly above chance even prior to SFC (Fig. 5a). However, social fear learning significantly enhanced the decoding accuracy, increasing from 57.4 ± 1.2% before SFC to 67.46% ± 1.57% decoding accuracy in the four trials 24 hours after SFC (mean ± SEM decoding accuracy across N=10 mice, Bonferroni multiple comparisons test: SE1&2 vs. SR1&2: p<0.01, SE1&2 vs, SR3&4: p<0.01). This is consistent with our single cell analysis and suggests that past experiences with conspecifics are incorporated into CA2 conspecific representations. In contrast, no significant change of decoding performance was observed for the Mock-SFC cohort, where the mean decoding accuracy plateaued at around 60%.

The increase in decoding accuracy following SFC is likely to be of behavioral significance as it was strongly correlated with both the amount of freezing behavior and the discrimination score for interactions with the CS+ and CS- during Social Recall trials 1 and 2 following SFC (Fig. 5b). Compared to trials 1 and 2, the relationship between decoding accuracy and discrimination score was weaker during trials 3 and 4, when there was no significant correlation between decoding and freezing behavior. The difference in correlations between the two pairs of trials likely reflects habituation or extinction of the fear response in the later trials. In contrast, we observed no significant correlation between behavioral read-outs and decoding accuracy for the Mock-SFC cohort during any stage of the paradigm.

The enhancement in decoding of CS+ from CS- could reflect a refinement in the representation of the CS+ alone, the CS- alone, or both. To distinguish among these possibilities we used the SVM classifier to decode interactions with the empty cup (during the empty cup recall trial) compared to when the same cups contained either the CS+ or CS- (in the social recall trials). We found that the accuracy of decoding both CS+ and CS- interactions relative to the empty cups was enhanced roughly equally after SFC, suggesting that both conspecific representations were refined by acquired valence (fear or safety) during memory formation (Fig. 5c).

Next we assessed if the threat-associated experience induced by SFC led to enhanced decoding of conspecific identity within a more naturalistic setting when both subject and stimulus mice were freely interacting. To this end, we exposed the subject mouse separately to the CS+ or the CS- when unconfined in an open arena during consecutive recall sessions 24 hours after SFC (Fig. 5d). We found that periods of CS+ and CS- interactions could be decoded successfully relative to periods when the subject was not engaged in social interactions, both when the subject mouse was approaching the stimulus mouse and when the stimulus mouse approached the subject mouse. Decoding accuracies were greater across all distances with respect to the CS+ as compared to the CS- (72.5 ± 1.61% for CS+ and 57.59 ± 1.87% for CS-; mean ± SEM decoding accuracy across N=5 mice, Bonferroni multiple comparisons test: Approach to CS+ vs. Approach to CS-: p<0.0001, Approach by CS+ vs. Approach by CS-: p<0.0001).

As CA2 encodes spatial as well as social information, we asked whether SFC also enhances the accuracy of spatial decoding or whether it was selective for social variables. To explore this question, we pooled CA2 activity data from interactions around the same cup location (ie, upper or lower cup) from two consecutive trials, irrespective of stimulus mouse identity (Extended Data Fig. 9). In line with previous work (Boyle, Posani et al., 2022), we found that CA2 activity allowed us to decode with above-chance level accuracy the spatial position of the social interaction. However, unlike social decoding, spatial decoding accuracy did not significantly increase after SFC and was not correlated with behavioral discrimination of the CS+ from CS- mice.

### CA2 population activity encodes a generalized or abstract representation of social valence irrespective of conspecific identity and location

Our data so far indicate that SFC enhances the ability of CA2 representations to discriminate the CS+ from CS- mouse. We next investigated whether this was solely based on a refinement of representations of the distinct social stimuli provided by the two mice or whether the CA2 representations incorporated the different valences associated with the mice. To assess if CA2 population activity contains generalized information on the valence associated with social encounters independent of social identity, subject mice underwent two SFC and recall sessions separated by one week, each with a different pair of CS+ and CS- stimulus mice (Fig. 6a). We focused on data from the first two trials of the recall session, with CS+ and CS- positions swapped between trials. We then determined whether a linear classifier could decode a given pair of CS+ and CS- mice when the classifier was trained on the other pair of CS+ and CS- mice. The ability of a classifier to accurately decode conditions not used in its training is termed its “cross-condition generalization performance” (CCGP), and provides a measure of whether a given brain region contains a generalized or abstract representation of a given variable (Bernardi et al., 2020).

Because we have two CS+ mice, two CS- mice, and two recall trials in which the mice are placed in cups at different locations, we have a total of 8 conditions, with a number of possible ways of assessing CCGP. To balance the binary classes for spatial location (up or down cup) and SFC session (week 1 or 2), we pooled CA2 activity around the CS- in the top and bottom cups in the two recall trials of week 1 to form one class and pooled CA2 activity around the CS+ in the two recall trials from week 2 to form the second class. We then trained a linear classifier to decode the CS+ and CS- from one set of conditions and measured the CCGP of the classifier by testing on the conditions not used for training, using pooled data for the two CS- trials from week 2 and pooled data for the CS+ from week 1. (Fig. 6b,c: “(1)”). We found that CA2 representations provided a high level of CCGP decoding accuracy that was greater than chance at a high level of statistical significance. Thus, we conclude that CA2 population activity provides a generalized represention of the valence of social interactions irrespective of social identity.

Next, we investigated if CA2 population activity permitted the decoding of social valence irrespective of *both* social identity and spatial position of the CS+ and the CS- mice (Fig. 6b,c: “(2)”). To this end, a linear decoder was trained on social interactions with a pair of CS+ and CS- mice around one cup location (eg, upper cup) and tested on a novel pair of CS+ and CS- in the remaining cup location (eg, lower cup). Again, the CCGP for valence was significantly above chance levels despite the differences in spatial and social identity conditions in the training and testing sessions. This indicates that CA2 population activity contains information on social valence which generalizes both across conspecific identity and spatial position. To determine whether CA2 also provides a generalized decoding of spatial position when the mice in the cups have an associated social valence, we trained the decoder to dissociate between interactions around the upper versus the lower cup when the cups contained stimulus mice of the same valence. We then tested the ability of the decoder to dissociate between social interactions around the upper versus lower cup with stimulus mice of the opposite valence (Fig. 6b,c: “(3)”). In line with previous work (Boyle, Posani et al., 2022), we found that CA2 population activity contains a representation of space independent of social valence.

High CCGP values using a linear classifier are a hallmark of low dimensional representations, in that a decoding hyperplane that separates one pair of conditions will also separate another pair of conditions for the same coding variable (Bernardi et al, 2020). In our case we find that we can decode social valence independent of spatial location of the CS+ and CS-. Similarly we can decode cup position independent of the valence of the mouse occupying the cup. Such low dimensional representations have a more limited ability to decode a range of variables (Bernardi et al., 2020; Boyle, Posani et al., 2022). One such variable is recall trial number; decoding trial number is referred to as the exclusive OR (XOR), as there is no commonality of spatial/social variables in the two classes when we group data by trial. Consistent with the idea that CA2 encodes valence in a low dimensional representation, we found that trial decoding accuracy was at chance levels (Fig. 6b,c: “(4)”).

## Discussion

Here, we provide for the first time a neural mechanism that contributes to the ability of an animal to discriminate a conspecific associated with threat from a conspecific associated with safety, based on a social fear conditioning paradigm. Moreover, we find that this discrimination requires the participation of hippocampal CA2 pyramidal neurons, which had previously been shown to be of critical importance for social novelty recognition memory. Importantly, the acquired fear responses following social fear conditioning were only manifest in the presence of a social context and not elicited by non-social cues (cup cages) present during conditioning, indicating the key role of social interactions in the conditioned fear response. To our knowledge these results provide the first evidence that the two major psychological components of social memory, the recognition of familiarity versus novelty and the recollection and discrimination of prior experiences with equally familiar individuals (Mandler, 1980), are mediated by a single class of neurons in a relatively small brain subregion.

Of further note, our study indicates that the dorsal CA2-dependent social specificity of social fear conditioning is dissociable from the association of threat with a general social context. Thus, silencing CA2 pyramidal neurons with chemo- or optogenetic methods did not prevent the acquisition or recall of social fear but rather prevented the discrimination of threat-associated from safety-associated individuals. Furthermore, CA2 plays a relatively selective role in social forms of fear conditioning. Thus, CA2 silencing did not impair the specificity of object fear conditioning, consistent with previous results that CA2 is not required for novel object recognition memory (Hitti and Siegelbaum, 2014; Oliva et al., 2020). Moreover, a previous study found CA2 silencing does not impair contextual fear conditioning in male mice (Hitti and Siegelbaum, 2014; although CA2 manipulations do modulate contextual fear memory in females; Alexander et al., 2019). These results provide a consistent picture that CA2 is of particular importance for encoding memory of social experiences, whether it be a simple assessment of social novelty versus familiarity or a more complex association of an individual with a particular aversive experience.

In principle, CA2 could participate in social fear specificity by providing information about social identity to downstream brain regions, such as the ventral CA1 region or the medial prefrontal cortex (Okuyama et al., 2016; Levy, Tamir et al., 2019; Biane et al., 2023), which then associate a given social identity with threat or safety. Alternatively, CA2 itself may participate in the association of a negative or positive valence with a specific individual’s identity. We found, based on calcium imaging of CA2 pyramidal neuron activity, that SFC strengthened the representations of both the CS+ and CS-, increasing the fraction of CA2 cells that were selectively activated during exploration of either stimulus mouse and enhancing the accuracy of the decoding by CA2 representations of the identities of the CS+ and CS-. Importantly, a linear classifier trained on one pair of CS+ and CS- mice could readily decode the CS+ from CS- using a different pair of social fear-conditioned mice. These results indicate that SFC leads to the incorporation of a general or abstract representation of social-threat versus social-safety associations into the CA2 representations of mouse identity.

How is the information on social valence and social identity contained in CA2 representations routed to downstream regions for the regulation of approach and avoidance behaviors? One likely route is through the connections from dorsal CA2 to ventral CA1, which projects to a number of brain regions important for different emotional and social behaviors, including the amygdala, a subcortical structure central for fear learning and fear behaviors (Fadok et al., 2018; Krabbe et al., 2018). A role for dorsal CA2 to ventral CA1 connections in social fear specificity is consistent with the finding that paired optogenetic activation of ventral CA1 social engram cells with a footshock can elicit a subsequent social fear response (Okuyama et al., 2016). Moreover, ventral CA1 (Okuyama et al., 2016) and its inputs from dorsal CA2 (Meira et al., 2018) are important for social novelty recognition memory. However, this form of memory depends on the outputs of ventral CA1 to the nucleus accumbens (Okuyama et al., 2016) and/or the medial prefrontal cortex (Phillips et al., 2019), rather than amygdala. Thus, dorsal CA2 may project to distinct subpopulations of ventral CA1 neurons that mediate social novelty and social fear behaviors through distinct output pathways, analogous to findings that distinct ventral CA1 subpopulations mediate social anxiety and contextual fear conditioning through separate output pathways to lateral hypothalamus and basal amygdala (Jimenez et al., 2018).

Our results on the encoding by CA2 of negative social valence extend previous work in head-fixed mice showing that the CA2 region encodes positive reward-associations valence during social odor-reward learning (Hassan, Bigler and Siegelbaum, 2023). These findings suggest that the same hippocampal subregion can encode both reward and aversive valence associated with conspecifics. Whether the same individual CA2 neurons encode both reward and social fear remains to be determined.

Associative learning processes, such as the generalization of conditioned fear to neutral social stimuli, have been implicated in maladaptive social fear (Lissek et al., 2008, 2014; Beckers et al., 2023; Gyles et al., 2023; Li et al., 2023) and have been shown to underlie social fear behaviors induced both by SFC and by social defeat (Xu et al., 2019; Ayash et al., 2020; Gyles et al., 2023). Neuropsychiatric disorders such as schizophrenia has been found to be highly comorbid with social anxiety disorders (Pallanti et al., 2004). Our finding that CA2 silencing leads to a loss of the selective display of safety and fear behaviors towards social stimuli, together with findings of alterations in CA2 properties in neuropsychiatric disorders in humans (Benes, 1998; Knable et al., 2004) and in mouse models of human disease (Piskorowski et al., 2016; Donegan et al., 2020; Modi et al., 2019), suggests a potential neural mechanism for social withdrawal and social fear generalization. Future research on CA2 may thus have important ramifications for understanding the etiology of and the development of potential treatments for social anxiety disorders and other disorders related to the aberrant assignment of valence to social stimuli (Nardou et al., 2023).

## Acknowledgements

We thank Larry Abbott, Daniel Salzmann, Lorenzo Posani, Sami Hassan and Robin Nguyen for critical discussions and comments on the manuscript, Chris de Solis for assistance with implementing preliminary SFC experiments, Anastasia Barnett for training of P.K. and members of the Siegelbaum lab for assistance throughout the work.

## Author contributions

P.K. and S.A.S. conceived of the project and wrote the manuscript. P.K. designed and implemented the modified SFC paradigm, implemented software, analyzed the behavioral and neural data and performed all surgeries and behavioral experiments except S.K.B. performed the viral injections for the optogenetic experiments and optogenetic fiber implant surgeries, D.M.G. performed the immunohistochemistry for the optogenetic experiment, N.S. performed the DeepLabCut analysis. Mouse and needle schematics in Figures 2a, 3a, 4a and EPM in extended data figures are from SciDraw, doi.org/10.5281/zenodo.3925901, doi.org/10.5281/zenodo.4912419, doi.org/10.5281/zenodo.3926205 and 10.5281/zenodo.5496320.

## Methods

### Animals

All experiments were approved by the Institutional Animal Care and Use Committee at Columbia University. Adult male and female mice (8-14 weeks old) were maintained on a 12-h light– dark cycle with ad libitum access to food and water. Subject mice used for all experiments were either from the C57BL/6J strain (Jackson laboratories; #000664) or transgenic mice on the C57Bl/6J background. Amigo2-Cre (Hitti and Siegelbaum, 2014) or Avpr1b-Cre (Jackson laboratories; #036876) mice and their wild-type littermates (all on a C57BL/6J background) were used for optogenetic and chemogenetic CA2 silencing and CA2 miniscope imaging experiments. No statistically significant differences between outcome variables were observed with respect to sex or between Amigo2-Cre and Avpr1b-Cre mice. Thus, data obtained from these cohorts were pooled. Stimulus mice were age-, sex- and weight-matched to subject mice and from the C57BL/6J or Avpr1b-Cre strains in experiments where CA2 was chemogenetically silenced in stimulus mice during social fear conditioning. Sample sizes were not predetermined, but are in line with those employed in the field. Experimenters were blind to animal genotype and mice were randomly assigned to a given group within the experimental conditions.

### Viral injections and surgical procedures

Mice were anesthetized with isoflurane and administered analgesics for all surgical procedures. Stereotaxic injections were performed using a nano-inject II (Drummond Scientific). The pipette was retracted 8 min after stereotaxic injection and after stereotaxic surgeries viruses were allowed to incubate for 3-4 weeks before behavioral testing.

#### Pharmacogenetic silencing of CA2

For the chemogenetic silencing of CA2 pyramidal neurons, Cre- dependent AAV2/5 hSyn.DIO.hM4D(Gi)-mCherry (Addgene; #44362) expressing the inhibitory hM4Di designer receptor exclusively activated by designer drugs (iDREADD) was injected bilaterally in the CA2 region of Amigo2-Cre or Avpr1b-Cre mice and wild-type littermate controls. A volume of 200 nL of virus (1.9×10^12^ pp/mL) was injected per hemisphere at AP: -2.0 mm, ML: +/-1.8 mm and DV: -1.5 mm from Bregma.

#### Optogenetic silencing of CA2

For optogenetic silencing of CA2 pyramidal neurons, AAV2/5-EF1a- DIO-hChR2-eYFP (1.9×1012 pp/mL) expressing the inhibitory opsin eArch3.0 was injected bilaterally in the CA2 region of Amigo2-Cre or Avpr1b-Cre mice and wild-type littermate controls. A volume of 200 nL of virus (1.9×10^12^ pp/mL) was injected per hemisphere at AP: -1.8 mm, ML: +/-2.1 mm and DV: -1.4 mm from Bregma. One week after injections, optical fiber assemblies (200 μm core, 0.37 NA, 3 mm, RWD Life Science Inc.) were implanted placed at AP: -1.8 mm, ML: +/-2.1 mm and DV: -1.2 mm from Bregma. Fibers were permanently fixed using dental cement.

#### Miniscope surgeries

A volume of 200 nL of pGP-AAV2/5-syn-FLEX-jGCaMP8m-WPRE (1.9×10^12^ pp/mL, Addgene; #162378) expressing the genetically encoded calcium indicator GcaMP8m was injected into the CA2 region of the right hemisphere at AP -2.0 mm, ML +1.8 mm, DV -1.7 mm from Bregma of Amigo2-Cre or Avpr1b-Cre mice (7-8 weeks old). Two to three weeks after stereotaxic injection, a Gradient Refractive Index lens (GRIN lens; Inscopix, 1.0 mm diameter, 4.0 mm height) was implanted. The skull was scored with a scalpel followed by a rectangular craniotomy (1.2×1.2 mm; centered at AP -2.0 mm, ML +2.2 mm). Following the craniotomy, the GRIN lens was lowered to a depth of DV -1.4 to -1.5 mm from Bregma at a 10° angle to position the lens surface parallel to the CA2 pyramidal layer. Finally, the lens was secured in place using Metabond dental cement. One week after the lens surgery, a baseplate was placed over the lens and secured with Metabond dental cement to enable rigid attachment of the miniscope for in vivo behavioral recordings.

### Immunohistochemistry

Mice were transcardially perfused using 0.9% saline followed by 4% PFA diluted in PBS. Brains were harvested and incubated in 4% PFA overnight and washed three times for 10 min in PBS the following day. 60-µm coronal sections were obtained with a Leica VT1000S vibratome and subsequently permeabilized and blocked for 2 h with 5% goat-serum and 0.5% Triton-X in PBS at room temperature. Sections were incubated overnight with primary antibodies, diluted in 5% goat-serum and 0.1% Triton-X in PBS, at 4°C. The following day, sections were washed three times for 10 min in PBS, followed by the application of secondary antibodies at room temperature for 4 h in 5% goat-serum and 0.1% Triton-X in PBS. Neuronal somata were visualized with a stain against the Nissl substance by using the NeuroTrace 435/455 fluorescent dye (1:200 dilution, Invitrogen; #N21479). All secondary antibodies were raised in goats, purchased from ThermoFisher Scientific and diluted at 1:500. Sections were mounted using fluoromount (Sigma-Aldrich) and images were acquired at 5x and 20x resolution using an inverted confocal microscope (Zeiss, LSM 700) and processed using FIJI software.

For eYFP and mCherry labeling, the first incubation was performed with chicken anti-GFP (1:1000, AVES Labs, #GFP-1020) and rabbit anti-RFP (1:1000, Rockland, #600-401-379). The secondary incubation was performed with anti-chicken conjugated to Alexa 488 (Thermofisher; #A11039) and anti-rabbit conjugated to Alexa 568 (Thermofisher; #A11011). For RGS14 and PCP4 labeling, the first incubation was performed with mouse IgG2a anti-RGS14 (1:50, UC Davis/NIH NeuroMab Facility, #73-170) and rabbit anti-PCP4 (1:400, Sigma-Aldrich; #HPA005792). The secondary incubation was performed with anti-mouse IgG2a or anti-rabbit conjugated to Alexa 488 (ThermoFisher; #A21131).

#### Analysis of viral expression

To confirm selective expression of the AAVs in CA2, the overlap between an expressed viral fluorescent marker and the CA2-specific markers RGS14 or PCP4 were assessed.

### Behavioral assays

Between all behavioral trials the arena and wire cups were cleaned with 70% alcohol wipes followed by cleaning with water and drying. Behavioral assays conducted with male versus female mice were conducted with at least a one week break in between.

#### Social Fear Conditioning (SFC) Paradigm

##### Apparatus

A modular shuttle chamber (51 cm length, 25 cm width, 30 cm height; Coulbourn, Harvard Biosciences) for rats with a grid floor for mice was employed for fear conditioning. The chamber was placed within an isolation cubicle for noise attenuation and connected to a precision shocker (Coulbourn H13-15 Precision Shocker). During recall, a rectangular arena (50 cm length, 25 cm width, 30 cm height; made in-house) was employed. Imaging Source DMK 37BUX252 video cameras mounted for top-down recordings of fear conditioning or recall sessions were employed for video recording at 30 Hz. Throughout all stages of the paradigm 3D printed cylinder-shaped plastic caps were placed on top of wire cage cups to prevent mice from climbing on the top of the cups.

##### Subject mice

Mice were habituated to the animal facility for at least one week after arrival from the vendor. Habituation to handling (5 minutes/day), the conditioning room (2 hours/day) and wire cage cups (2 hours/day) took place for two consecutive days before the start of the behavioral tests. Mice were randomly assigned to the fear-conditioned experimental group or the unconditioned control group, with the same behavioral paradigm being followed for the latter with the exception of foot shock delivery. Subject mice were singly housed beginning two hours prior to the fear conditioning stage of the experiment to allow for habituation to the homecage and maintained singly housed throughout the behavioral paradigm to prevent social fear extinction due to exposure to social stimuli. Habituation, fear conditioning and recall sessions took place during the same time of the day during the light phase.

##### Stimulus mice

Stimulus mice went through the same habituation routine as subject mice, but were additionally habituated to being placed under wire cage cups twice for 5 min on each habituation day. Stimulus mice were singly housed for one week prior to the social fear conditioning and throughout the paradigm to prevent odor cross-contamination with other mice. Two animals from distinct litters from the subject mice served as the social stimulus mice. The same two stimulus mice were typically used for a cohort of around 10 subject mice and designated at random as either the CS+ or the CS- for each subject mouse. Stimulus mice and control mice were housed in distinct rooms within Columbia University’s animal facility and kept separate throughout the entire experiment except for fear conditioning and recall phases. Stimulus mice were placed on plastic bases to prevent them from receiving foot shocks.

##### Social fear conditioning

Fear conditioning consisted of two phases of 5 minutes each. During a non-social ‘Cups’ stage mice were allowed to freely explore for 5 min the fear conditioning chamber containing two empty wire cage cups placed on opposite sides of the chamber. During the social fear conditioning stage both wire cage cups were replaced by identical wire cage cups containing the stimulus mice. The two stimulus mice were randomly assigned as the CS+ or the CS- for each experimental mouse and placed at random on either the right or left side of the fear conditioning chamber. Subject mice that were conditioned received a foot shock (1 second duration, 0.7 mA pulsed DC current), which was manually delivered whenever a subject mouse was judged to interact with the CS+ mouse, based on the criteria that the head of the subject mouse was both inside a social interaction zone, defined as an area within 2.5 cm of the cup containing the CS+ mouse, and oriented towards the center of the CS+ cup. The fear conditioning stage ended either after 5 minutes of explorations or when 4 foot-shocks had been administered, at which time the mice were returned to their homecage.

##### Social fear recall

Social fear recall trials were performed 24 hours after fear conditioning in the same behavioral room but in a novel arena, rotated by 90 degrees with respect to the experimenter, with different lighting (red LED light strips) than the fear conditioning arena and with smooth acrylic flooring in place of the metal grid. Similar to the fear conditioning trial, a recall trial consisted of two phases of 5 minutes each. Subject mice were exposed to a non-social ‘Cups’ recall stage, where mice were allowed to freely explore two empty wire cage cups placed on opposite sides of the arena. This was followed after approximately 2 min by a 5 min ‘Social’ recall stage, where the subject mouse was exposed to the CS+ and CS- stimulus mice previously encountered during fear conditioning. CS+ and CS- positions (distal versus proximal parts of the arena, relative to the experimenter) were assigned at random for the first trial of the recall session. Their positions were then swapped for every consecutive trial with different subject mice.

##### SFC behavioral paradigm for miniscope recordings

Prior to calcium imaging sessions using a miniature head-mounted microscope (miniscope), mice were habituated with a dummy miniscope until comfortable. A commutator was employed during all recording sessions to enable rotation of the miniscope cable, thus allowing mice to move freely in the arena. Prior to behavioral testing, mice were allowed to acclimate to the behavioral room for 2 hours. An nVista 3.0 Inscopix miniaturized microscope was inserted into the baseplate and used to record fluorescence signals from dorsal CA2 pyramidal neurons expressing GcaMP8m during free behaviors using Inscopix data acquisition software (30 frames per second, 50-ms exposure, 0.2-0.4 mW/mm^2^ illumination intensity). The focal plane was optimized for each mouse individually and maintained throughout all recording sessions. To align behavior and calcium videos, a 5-V TTL pulse was triggered from an AMi-2 Optogenetic interface at the start of calcium recordings through Anymaze software synchronous with behavioral video recordings, both acquired at a rate of 30 Hz. Habituation to room and handling was conducted as described for the SFC behavioral paradigm.

Miniscope recording experiments were performed in two behavioral paradigms, a Mock SFC paradigm and an SFC paradigm, separated by one week. In the Mock SFC paradigm, mice were allowed to explore for 5 min a recall arena containing two empty wire cage cups on opposite sides of the arena. Two novel stimulus mice were then placed under the cups and the subject mouse was allowed to explore the arena for two additional 5-min trials, with the position of the stimulus mice swapped between the two trials. Mice were then returned to their home cage for 2 hours, after which they were allowed to explore the fear conditioning chamber with two empty cups for 5 minutes, followed by an additional exploration trial in which the same two stimulus mice were present in the cups, with positions randomly assigned to right or left cup. No shocks were delivered throughout the sessions. Mice were then returned to their home cage for 24 hours and then reintroduced to the recall arena for a 5 min trial of exploration of the empty cups ( ‘Cups’ stage). This was followed by four consecutive 5 min recall trials in which the same stimulus mice encountered previous day were present in the cups (‘Social’ stage). Trials were separated approximately by 2 min where the arena was cleaned and positions of the stimulus mice were swapped in each consecutive trial. After one week, mice underwent the SFC paradigm. The paradigm was similar to the Mock SFC paradigm except that subject mice were administered foot shocks during exploration of the designated CS+ stimulus mouse in the fear conditioning chamber, as described above during social fear conditioning. The positions of the stimulus mice were chosen at random for the first recall session and swapped for the following three recall sessions. The arena was quickly cleaned both prior to and between the recall session with the general cleaning procedure described above.

##### Social memory assay

Subject mice were habituated for 20 min to a rectangular arena with two empty wire cups (radius 5 cm) on opposite sides for four days. After habituation on day four, two novel stimulus mice were placed under the wire cups. In learning trials, the subject mouse was allowed to explore the arena with the novel stimulus mice for 5 min followed by another 5 min of exploration with the position of the stimulus mice swapped. The subject mouse was then placed into a holding cage, followed by re-introduction to the same arena 30 min later for a 5 min recall trial in which the arena again contained two cups, each with a stimulus mouse. One of the cups contained one of the now familiar stimulus mice encountered in the learning trials (F), chosen at random, and the other cup contained a third novel stimulus mouse (N). Social memory is manifest by the innate preference of the subject mouse to explore the more novel of the two stimulus mice. Social interaction (SI) was defined as times when the subject’s head was oriented towards the center of a cup containing a stimulus mouse within a 7.5 cm radius of the cup center (1.5x the radius of the cup; social interaction zone). We measured the total SI time during the recall trial for mouse N and mouse F and used this to calculate the following social discrimination index (DI) as a measure of social memory:

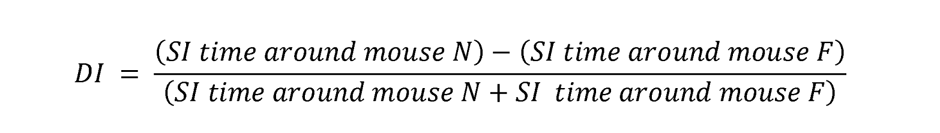

##### Elevated plus maze

Mice were transferred to the behavioral testing room 2 hours before the experiment for habituation to the room. For testing, mice were placed in the center of an elevated plus maze (Harvard Apparatus; two enclosed arms with 15-cm high walls and two open arms, arms 65 cm long and 6 cm wide; maze elevated 40 cm from the floor) facing the open arm of the maze and away from the experimenter. Mice were allowed to explore the maze for 8 min. The position of the head was tracked with DeepLabCut to calculate the number of entries into each arm, the time spent in each arm and the distance traveled.

##### Object Fear Conditioning (OFC) Paradigm

We performed an object fear conditioning (OFC). Experiment closely based on the SFC protocol, except we used two novel objects in place of the novel stimulus mice and the objects were not placed under cups. Different objects made of plastic or rubber (all about 10 cm tall and 5 cm × 5 cm wide) were used for the OFC paradigm and randomly assigned as the CS+ or the CS- for each experimental mouse. Both fear conditioning and recall stages were preceded by 5 minutes of exposure to the respective empty arena.

##### Optogenetic CA2 silencing during social fear encoding or recall

Mice were habituated for four days to the optogenetic apparatus by being allowed to freely explore the fear conditioning or the recall arena for 20 min while being connected to fiber optic leads attached to an optic rotary joint (Doric Lenses) centered above the arena. For photostimulation of eArch3.0, ferrules were connected to a 532- nm laser diode (OEM Laser Systems). A Master-8 pulse stimulator (A.M.P.I.) was used to control the laser, which was adjusted to around 8 mW for somatic stimulation using a digital power meter console (ThorLabs). Green light pulses were delivered continuously during the 5 min SFC encoding trial to inhibit CA2 pyramidal neurons by activating eArch3.0 bilaterally expressed in CA2 pyramidal neurons of Amigo2-Cre or Avpr1b-Cre mice. We used independent cohorts of mice to silence CA2 during encoding or recall. Control wild-type littermates injected with the same Cre-dependent virus received identical light pulses.

##### Chemogenetic silencing of CA2 during behavioral assays

Three to four weeks after stereotaxic viral injections of AAV2/5 hSyn.DIO.hM4D(Gi)-mCherry, Amigo2-Cre, Avpr1b-Cre or wild-type littermates were habituated to scruffing and IP saline injections for four consecutive days. Mice were then injected 30 minutes prior to behavioral experiments intraperitoneally with 5 mg/kg of the iDREADD agonist clozapine-n-oxide (CNO; Cayman Chemical, #<u>34233-69-7</u>) and returned to their homecage until behavioral testing. The volume of saline and CNO injections were matched between habituation and testing days.

##### Chemogenetic silencing of CA2 in stimulus mice during behavioral assays

Social memory tests and SFC paradigms where CA2 was silenced in Avpr1b-Cre^+^ stimulus mice were performed approximately 3-4 weeks after stereotaxic viral injections of AAV2/5 hSyn.DIO.hM4D(Gi)-mCherry in the dorsal CA2 of the stimulus mice. Social memory tests were performed with intraperitoneal administration of CNO to the stimulus mice 30 min prior to the first social memory encoding session. One week later, a social memory test was conducted with the same stimulus mice and a novel cohort of subject mice, whereas intraperitoneal saline injections were administered to the stimulus mice 30 mins prior to the first social memory encoding session. For the SFC paradigm with CA2-silenced stimulus mice (i.e., the CS+ and the CS-), stimulus mice were administered intraperitoneal CNO 30 min before social fear conditioning (day 1 of the SFC paradigm). The same pair of CS+ and CS- mice received one week later intraperitoneal saline injection 30 min before social fear conditioning and was employed for a novel cohort of subject mice. Both the social memory and the SFC assays with CA2- silenced stimulus mice were in addition conducted with a second pair of stimulus mice and independent cohorts of subject mice.

### Behavioral analysis

#### Pose tracking with DeepLabCut

DeepLabCut software (v.2.3; Mathis et al., 2018) based on ResNet- 50 was used for markerless pose estimation of social fear conditioned and unconditioned control mice during the ‘Cups’ and the ‘Social’ stages of recall. DeepLabCut was trained based on the following 9 markers on the body of the subject mouse: (1) Tip of the nose, (2) left ear, (3) right ear, (4), center of head (between both ears), (5) left side of waist, (6) right side of waist, (7) body center, (8) base of the tail, (9) center of the tail, and (10) tip of the tai and 4 additional markers indicating the corners of the arena, as well as two markers indicating the center of the wire cage cups placed on opposite sides of the arena. The training set was composed of 320 videos and a total of 2,000 labeled frames randomly sampled with the built-in k-means algorithm and run for 500,000 iterations. Coordinates predicted with a likelihood below 95% were replaced by interpolation. Manual labeling of markers was done with the experimenter blind to genotype and experimental condition.

#### Head direction analysis

The head direction of mice was calculated based on the vector connecting the nose and body center. A subject mouse was taken as being oriented towards a stimulus mouse when the center of the relevant wire cage cup was within 20° to either side of this vector. Social interactions (SI) were defined as times when the subject’s head was oriented towards the center of a cup containing a stimulus mouse within a 7.5 cm radius from the cup center (1.5x radius of the cup radius).

#### Manual annotation of videos

Video labels for supervised classification were obtained through manual annotation with the BORIS software (Friard and Gamba, 2016), based on top-down video recordings obtained from the ‘Social’ stage of recall trials from non-conditioned controls, social fear conditioned mice and social fear conditioned mice during chemogenetic silencing trials. Behaviors were categorized as (1) “Freezing” (no movement except for respiration and heartbeat for at least 1 s), (2) “Approach” (approach to a stimulus mouse in stretched posture), (3) “Avoidance” (rapid retreat from a stimulus mouse), (4) “tail rattling” (fast waving movement of the tail), (5) “Social interaction” (i.e., within social interaction zone and head oriented towards cup center) or (6) “Other” for any other type of behavior. A total of 160 videos with a framerate of 30 Hz were annotated, resulting in a total number of 1,440,000 annotated frames. All annotations were carried out with the experimenter blind to the genotype and experimental condition.

#### Feature extraction for supervised and unsupervised classification of behaviors

The following behavioral features were used to classify behaviors of the subject mice both in the presence and absence of social stimulus animals: (1) Distance of body center to cup centers; (2) Distance of head to cup centers; (3) Distance of nose to cup centers; (4-7) Distance of the head to each of four corners; (8- 11) Distance of head to each of four walls; (12) Distance between body center and nose; (13) Distance between the left and right waist; (14) Speed of the subject mouse (running average over 2 frames); (15) Head orientation of the subject mouse with respect to cup centers; (16,17) all x,y coordinates for the relevant markers as obtained with DLC.

#### Feature preprocessing

Behavioral features for supervised and unsupervised classification were normalized to the [0,1] range with Python’s sklearn.preprocessing StandardScaler to account for variability in animal size and slight differences in camera position. Features with values outside the 1st and 99th percentile were removed to exclude outliers.

#### Supervised classification of behaviors with a long short-term memory (LSTM) recurrent neural network

To classify behaviors into the six classes as described above, a long short-term memory (LSTM) recurrent neural network was implemented using Python’s TensorFlow package to account for the sequential nature of behaviors. Manual annotations were used as the ground truth and prediction relied on the 16 behavioral features defined above. The LSTM model architecture comprised an input layer, the LSTM layer with 100 memory units and a fully connected softmax layer enabling multiclass classification. Custom python code was implemented to generate sequences of 90 frames (3 s) with a shift of at least ten frames between consecutive sequences. Data was stratified such that train, validation and test sets contained sequences extracted from different segments of the video sets to minimize dependency. The Adam algorithm for gradient descent was used for optimization of the model together with the categorical cross-entropy loss function, and classification accuracy was employed as the outcome metric. A dropout and recursive dropout of 0.2 were used for regularization. Shuffled data sets were generated by random permutation of the behavior labels. Python’s scikit-learn kernel density estimation function was then used for visualizing the distribution of predicted behaviors.

### Analysis of one-photon calcium imaging data

#### Preprocessing of one-photon calcium data

Videos of calcium-dependent fluorescence signals from separate sessions within one day were concatenated within Inscopix Data Processing Software (IDPS) as ‘Time Series’. Time Series were corrected for defective pixels and 4x spatially down-sampled. Background fluorescence was removed using a spatial band-pass filter based on default parameter values and fluorescence videos were motion-corrected to the mean image using the Inscopix motion correction algorithm. Cell identification was implemented with the Inscopix CaImAn implementation with the average cell diameter evaluated with the Inscopix measurement tool, with a minimum pixel correlation parameter of 0.9, a merging factor of 0.9 and a minimum peak-to-noise ratio of 15. Default values were used for all other parameters and the temporal traces were normalized to dF/F, i.e., the change of a cell’s fluorescence normalized to the mean fluorescence across the entire recording.

#### Single cell analysis of calcium data

Calcium signals were measured in 100 ms bins during periods when the subject mouse was in an interaction zone around a cup (either empty or containing a stimulus mouse) and oriented towards the center of a given cup. To determine whether the activity of a given neuron was modulated by social interactions, we compared the calcium signals when the subject mouse explored the CS+ or CS- to the calcium signals recorded during the same trial when the subject mouse was not engaged in social interactions. We used a non-parametric Wilcoxon rank-sum test to determine whether there was a significant difference in calcium signals during social compared to non-social exploration. Cells whose activity during social exploration with the same stimulus mouse differed significantly from baseline activity (pL≤L0.01) during two consecutive social recall trials and across both spatial positions (upper and lower cup) were classified as CS+ and/or CS- selective, according to the nature of the responses. Cells which were both CS+ and CS- selective were classified as ‘mixed selective’. Cells whose activity during social explorations was significantly different from baseline across one spatial position independent of stimulus mouse identity were classified as spatially selective.

#### Linear population decoding

We used a support vector machine (SVM) with a linear kernel to decode calcium responses using a Python implementation of the libsvm package (libsvm3.23; Chang and Lin, 2011). We trained the decoder on a subset of 90% of the data and then tested the performance on a subset of 10% of data that was withheld from the training data set. Features were constructed by averaging the DF/F calcium fluorescence signals of each cell over 100 ms bins, and by assigning labels to the samples according to the mouse’s behavior (e.g., social interaction with the CS+ or the CS-). Sample sizes were balanced by class and across mice. Training, validation and test sets were stratified such as to contain samples from distinct social interaction periods. To decode information on social identity of the CS+ versus the CS+ independently of spatial location based on CA2 calcium activity, binary classification was performed on data combined from two consecutive trials, where the positions of the CS+ and the CS- are swapped, with time spent exploring right and left cups balanced. To decode information on spatial position based on CA2 calcium activity, binary classification was used with calcium data combined from social interactions around the upper versus the lower interaction zones of the arena, irrespective of the interaction partner (CS+ or CS-), with time spent exploring the two stimulus mice balanced. The binary SVM classifier was then employed to decode spatial position (upper versus lower interaction zone). Decoding for the open arena assay was implemented with the same scheme, except that binary classes were constructed based on instances where subject and stimulus mice were oriented towards each other vs. when they were not, matched for distance and velocity of the moving agent. In addition, classes were balanced for the four quadrants of the recall arena.

#### Cross-condition generalization performance

To calculate the cross-condition generalization performance (CCGP), a linear SVM was trained to dissociate a dichotomy under one set of conditions and tested on the same dichotomoy on data from a different set of conditions. For example, to determine the CCGP for the dichotomy valence, we trained the classifier based on interactions with the CS1+ versus CS2- and then tested that classifier for its ability to decode the interactions with the CS2+ versus CS1-. We then reversed training and test data sets and measured CCGP as the average decoding performance for the two determinations. For cross-identity and cross-position social valence CCGP (Fig. 5b, “(2)”) and cross-identity position CCGP (Fig. 5b, “(3)”), training and test sets were constructed from data pairs with opposing non-decoding variables. For instance, when decoding cross-identity *and* cross-position social valence CCGP, one training set contained data from social interactions around the upper cup while one test set contained left-out data from social interactions with a different pair of CS+ and CS- around the lower cup. The CCGP decoding was repeated with swapped training and test sets and for (2) and (3) with swapped non-decoding variable associations. The reported CCGP was calculated as the average decoding accuracy for all relevant training and testing schemes whereby sample sizes were balanced across all possible classes. The CCGP null-model was estimated by randomly shuffling neuron indices, as described in Boyle, Posani et al., 2022.

#### Generalization of decoder performance across social fear conditioning

To test the hypothesis that the increased separability of CS+ and CS- representations was driven by changes in the CS+ representation, two distinct linear SVMs were trained either to decode social interactions with the CS+ or the CS- from baseline activity prior to social fear conditioning. Then, each classifier was tested on calcium data acquired during the same type of behavior after fear conditioning, from calcium data obtained from cells registered across the two days.

#### Permutation testing and empirical chance level calculation

Empirical chance level was calculated by randomization of classification labels and empirical p-values were calculated as the fraction of shuffled classification accuracies above the accuracies obtained based on the ground truth.

### Statistical analysis

All statistical analyses were implemented in either Python (v.3.11.1) or R (4.2.2). Non-parametric statistical tests comprised the Wilcoxon rank-sum test, Wilcoxon signed-rank test, *t*-test, and the Kruskal–Wallis test as appropriate. One-way analysis of variance (ANOVA), two-way repeated-measures ANOVA, and t-tests were used where indicated. t-tests were two-sided except where indicated otherwise. Midlines of box plots indicate median, while the range of the box plot indicates the upper (75%) and lower (25%) quartiles. Box plot whiskers include data within a 1.5x interquartile range. A significance threshold of αL=L0.05 was employed except when indicated differently, with NS, not significant thus indicating *P*L>L0.05 and the following used otherwise: *pL<L0.05, **pL<L0.01, ***pL<L0.001, ****p < 0.0001. All behavioral, imaging and optogenetics experiments were replicated in multiple subject animals with similar results.

### Python packages

The majority of the code was implemented in Python, and makes use of the following packages: ‘os’, ‘sys’, ‘utils’ and ‘re’ for system administration; ‘numpy’, ‘scipy’ and ‘pandas’ for data processing, data analysis and general data handling; ‘scikit-learn’ for preprocessing of the data for classification and k-means clustering; ‘TensorFlow’ and ‘libsvm’ for LSTM and Support Vector Machine classification respectively; ‘Matplotlib’, ‘seaborn’ and ‘plotly’ for data visualization. Plotly data visualization was used for polar plots based on modified custom code from (Ashaber, Tomina, Kassraian et al., 2021).

---

**Extended Data Figure 1:**
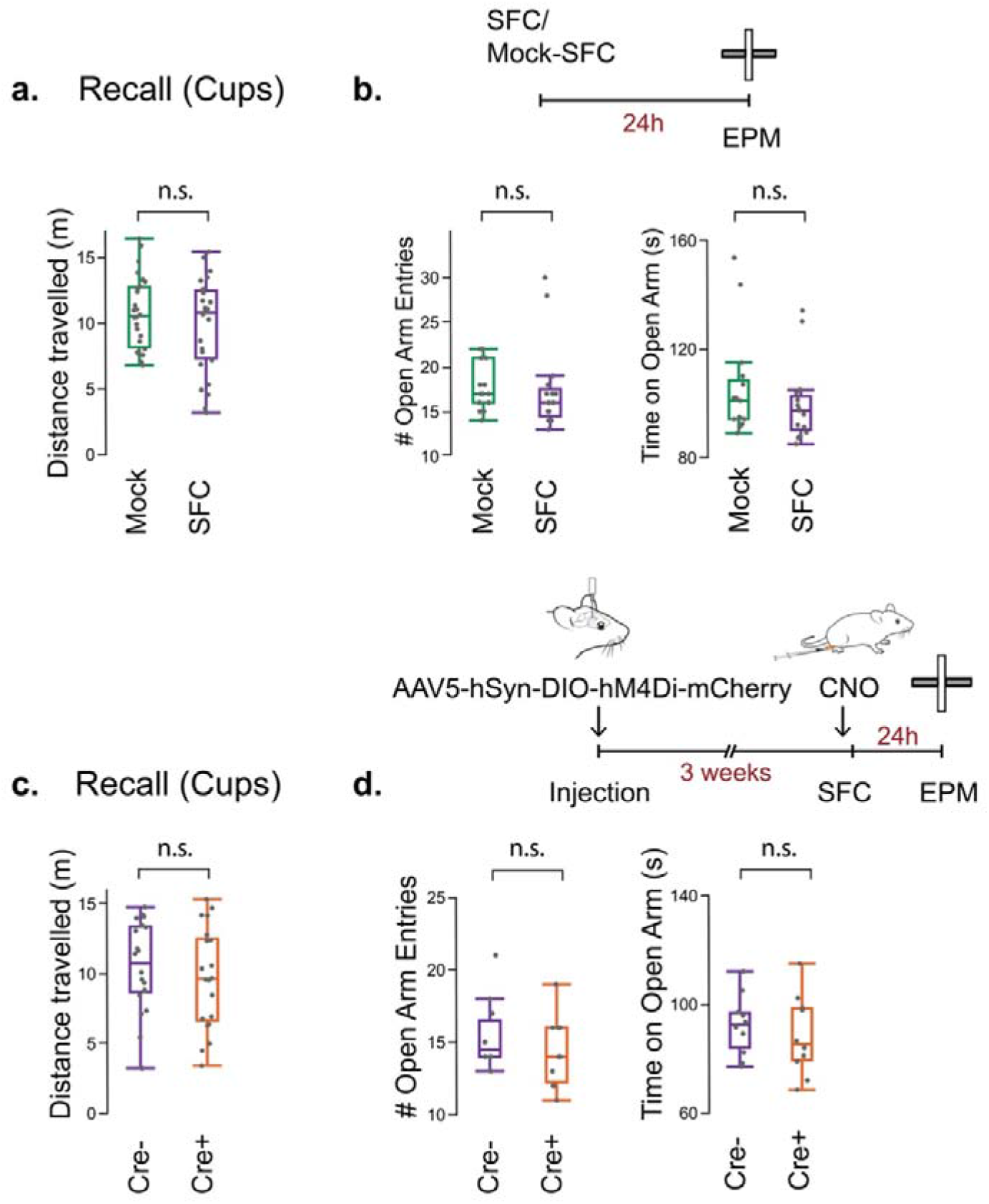
Assessment of general anxiety-like behavior and locomotor activity. **a.** No significant differences between Mock-SFC and SFC cohorts are observed for the distance traveled during the non-social ‘Cups’ recall trial 24 hours after SFC. Unpaired two-sample t-test: Distance traveled t=1.019, p=0.313. N=13 female mice plus N=13 male mice per cohort. **b.** No significant differences between Mock-SFC and SFC cohorts are observed during exploration of the Elevated Plus Maze (EPM) 24 hours after SFC. Unpaired two-sample t-test: Open arm entries t=0.32, p=0.751, Time on open arm: t=0.934, p=0.358. N = 8 females plus 7 males per cohort. **c.** No significant differences between SFC Cre- and Cre+ cohorts with CNO injection 30 mins prior to the SFC session are observed for the distance traveled during the non-social ‘Cups’ recall trial 24 hours after SFC. Unpaired two-sample t-test: Distance traveled t=0.767, p=0.447. N=10 female mice plus N=10 male mice per cohort. Avpr1b-Cre+ and Amigo2-Cre+ numbers balanced across cohort. **d.** No significant statistical differences between SFC Cre- and Cre+ cohorts are observed during exploration of the EPM 24 hours after SFC, with CNO administration 30 min prior to the EPM assay. Unpaired two-sample t-test: Open arm entries t=1.092, p=0.289, Time on open arm: t=0.637, p=0.531. N = 5 females plus 5 males per cohort, Avpr1b-Cre+ and Amigo2-Cre+ numbers approximately balanced across cohort. No significant effect of sex observed for a., b. and no significant effect of sex and between Cre+ and Cre- cohorts observed for c.,d..

**Extended Data Figure 2:**
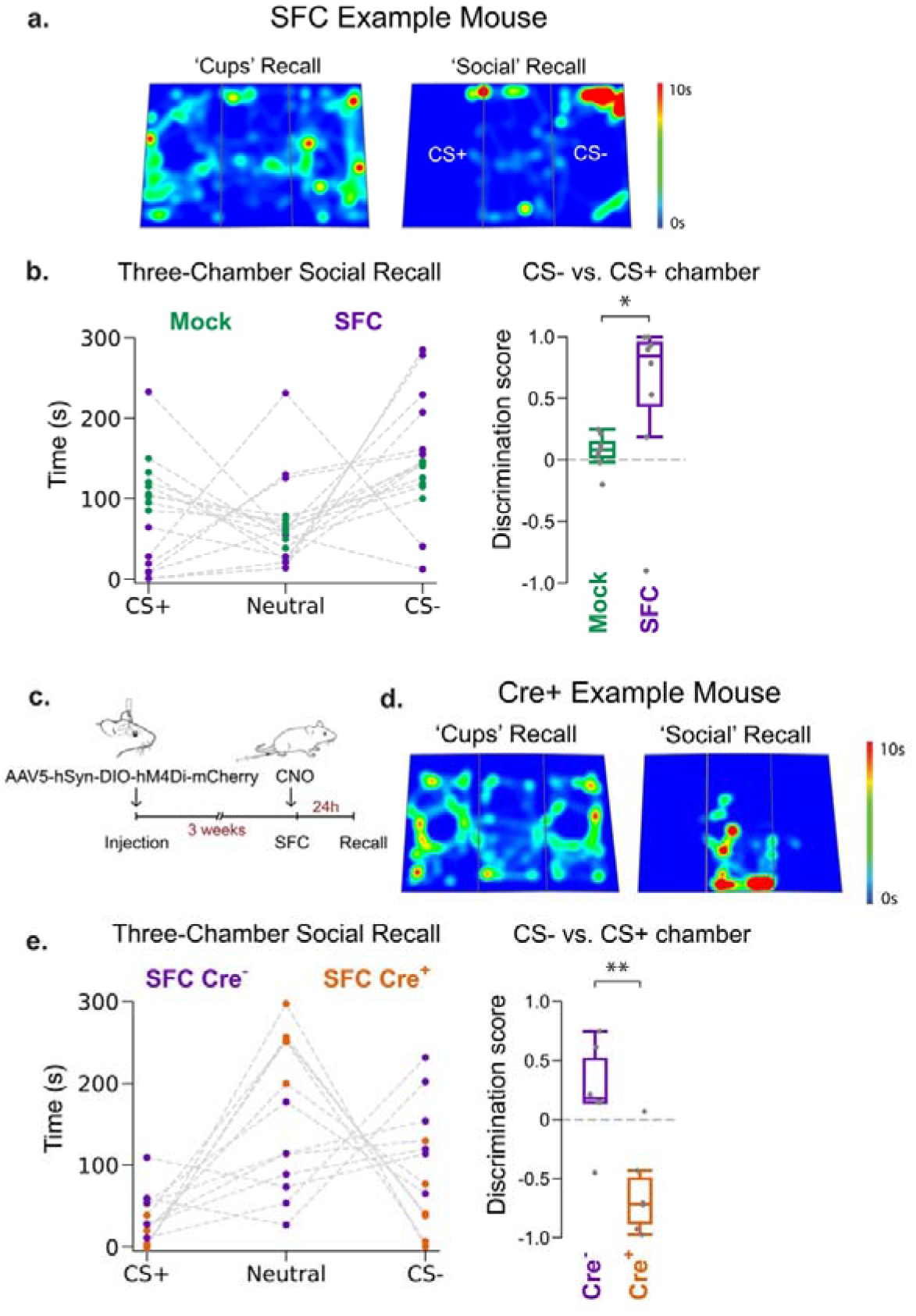
Social fear recall in the Three-Chamber Arena. **a.** Example heatmap during recall in a three-chamber arena for one SFC mouse, illustrating its preference for the CS- chamber. **b.** Left: Time spent in each chamber of the three-chamber recall arena for Mock-SFC and SFC cohorts. Right: The SFC cohort shows a significantly greater preference for the CS- chamber than the Mock-SFC cohort. Unpaired two-sample t-test: Discrimination score t=2.448, * p=0.039. N= 4 males plus 4 females per cohort. **c.** Approximately 3-4 weeks after Cre-dependent AAV2/5 hSyn.DIO.hM4D(Gi)-mCherry injection in CA2, Cre+ mice and Cre- controls were intraperitoneally injected with CNO 30 min prior to SFC. **d.** Example heatmap for one SFC Cre+ mouse during recall trials (no CNO), showing its non-selective avoidance of CS+ and CS- and retreat to the neutral middle chamber. **e.** Left: Time spent in each chamber of the three-chamber recall arena 24 hrs after SFC for Cre- and Cre+ cohorts. Right: The SFC cohort shows a significantly greater preference for the CS- versus the neutral chamber than did the Cre+ cohort. Unpaired two-sample t-test: Discrimination score t=3.647, ** p=0.0044. N = 4 female mice plus N = 4 male mice per cohort. Avpr1b-Cre+ and Amigo2-Cre+ numbers balanced across cohort and sex with no significant effect of Avpr1b-Cre+ versus Amigo2-Cre+ genotype and sex observed.

**Extended Data Figure 3:**
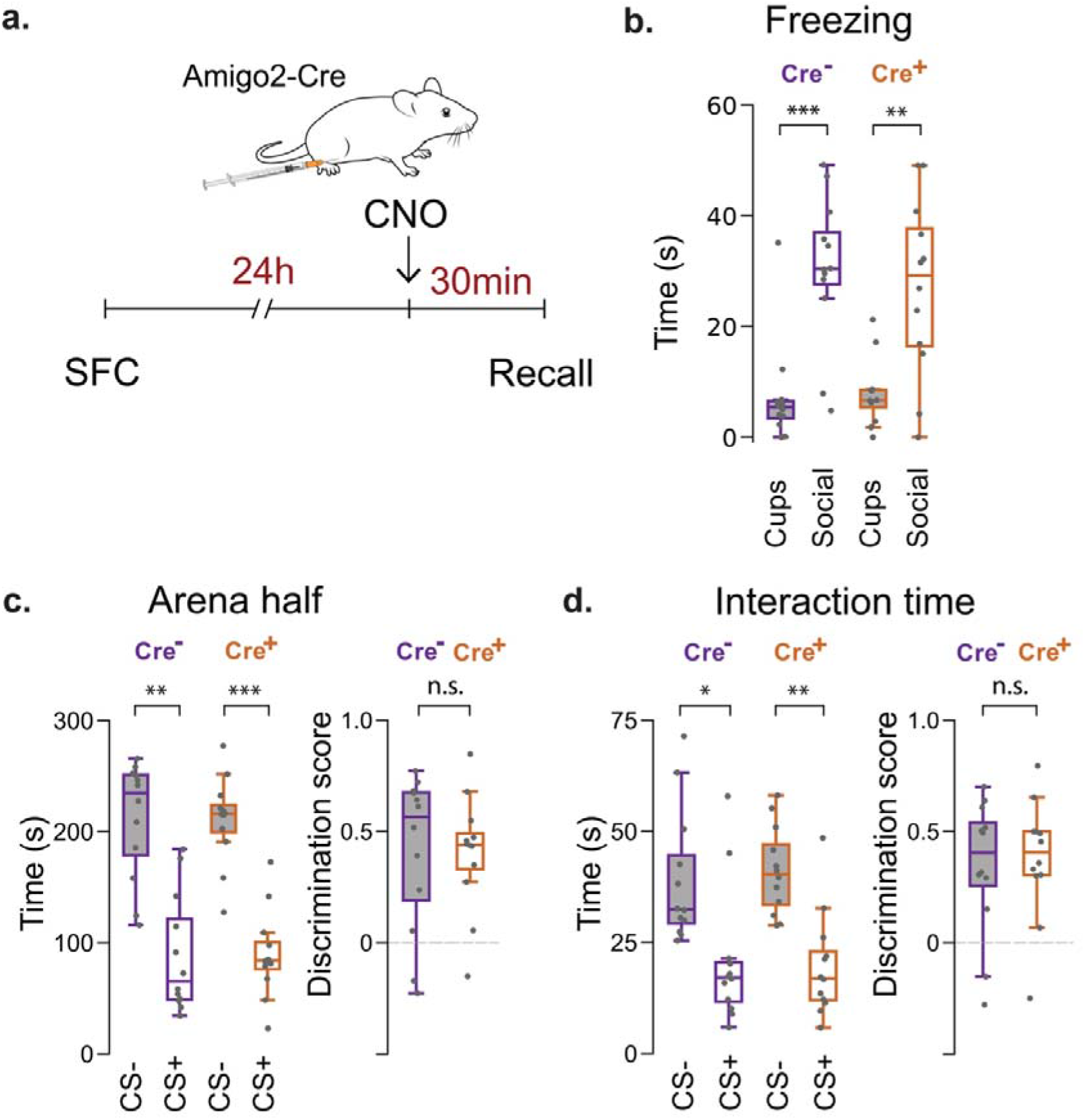
Assessment of genotype effects on SFC responses. **a.** Experimental timeline: To assess whether SFC responses differ in Amigo2-Cre+ versus Amigo2-Cre- littermates in the absence of iDREADD expression, Cre- and Cre+ littermate controls undergo SFC on experimental day 1. 24 hours later, mice are intraperitoneally injected with CNO 30 min prior to recall. **b.** SFC Cre- and Cre+ cohorts display similar freezing levels during the non-social ‘Cups’ recall trial and the Social recall trial (two-way repeated measures ANOVA: Cohort x Stage, F(1,44) = 0.871, p = 0.68), with both displaying significantly increased time freezing during the Social recall trial. **c.** Both Cre- and Cre+ cohorts spend a significantly greater time in the CS- versus the CS+ half of the arena, with no significant difference between genotypes (two-way ANOVA: Cohort x Arena half, F(1,44) = 1.97, p = 0.52). Discrimination score based on time in arena half, unpaired t-test t = -0.033, p = 0.973. **d.** Cre- and Cre+ cohorts interact significantly more with the CS- than the CS+ with no difference between genotypes (two-way ANOVA: Cohort x Stimulus mouse, F(1,44) = 1.28, p = 0.47). Discrimination score based on interaction time, unpaired t-test: t = 0.292, p = 0.772. Bonferroni post-hoc tests: * p < 0.05, ** p<0.01, *** p<0.001. N = 6 female mice plus N=6 male mice per cohort, Amigo2-Cre+ mice with sex balanced across cohorts. No significant effect of sex observed.

**Extended Data Figure 4:**
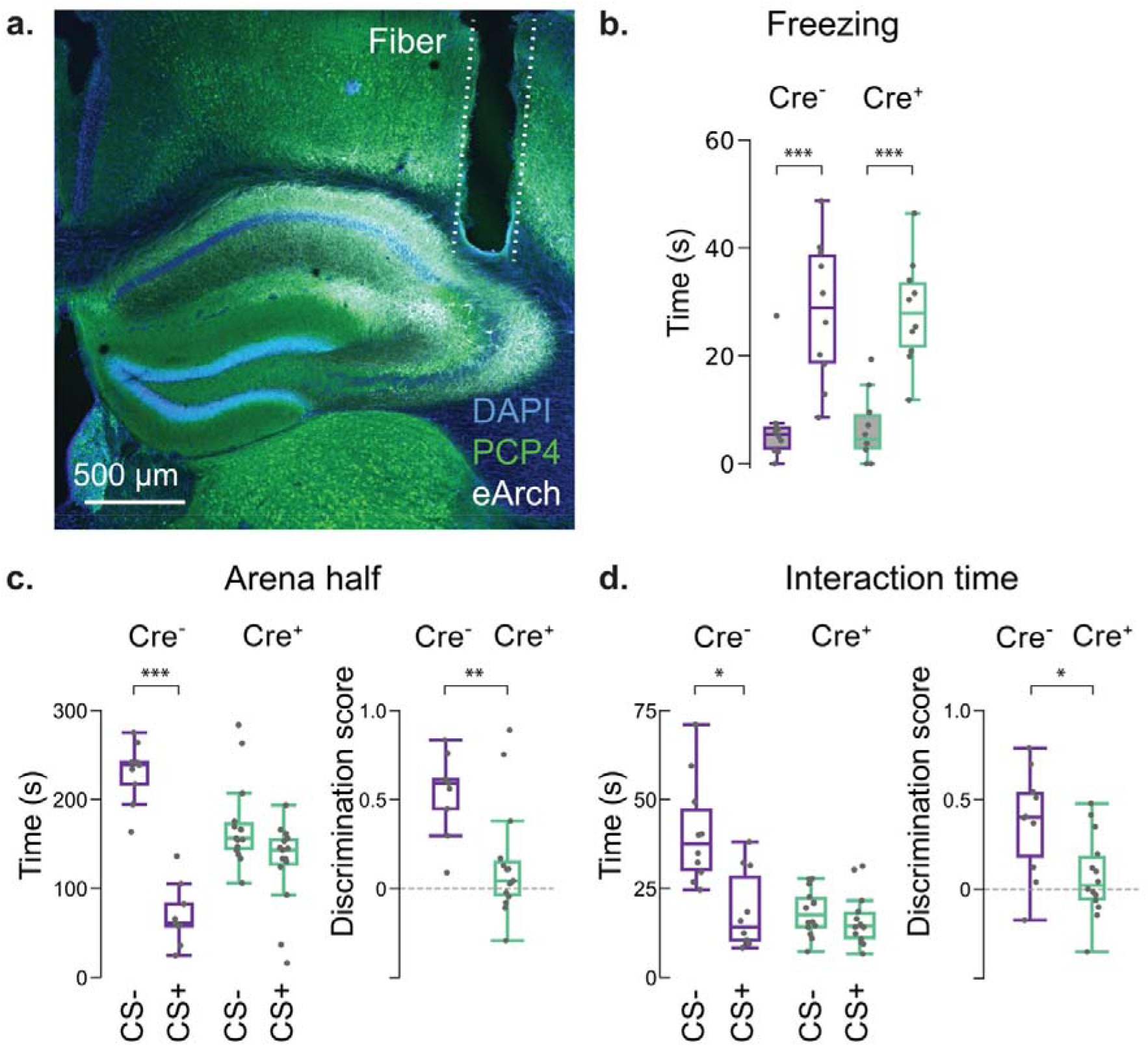
Optogenetic silencing of CA2 pyramidal cells during social fear conditioning disrupts social fear specificity. **a.** For optogenetic silencing of CA2 pyramidal cells during SFC (day 1), Amigo2-Cre (Cre+) mice and wild-type littermates (Cre-) were injected in CA2 with AAV-DIOeArch3.0-eYFP and implanted with bilateral ferrules for optic fiber probes above CA2. Example confocal image from an Amigo2-Cre mouse showing optical fiber tract and eArch3 expression in CA2, defined by co-expression of CA2 marker PCP4. CA2 was continually illuminated in Cre+ and Cre- groups with yellow light during 5-min SFC learning trial. **b.** SFC Cre- and Cre+ cohorts display similar freezing levels during the non-social ‘Cups’ recall trial and the Social recall trial (two-way repeated measures ANOVA: Cohort x Stage, F(1,40) = 0.75, p = 0.72), with both displaying significantly increased time freezing during the Social recall trial. **c.** Cre- cohorts show a preference for exploring arena half containing CS- whereas Cre+ cohorts spend comparable times in the CS+ and CS- arena halves (two-way ANOVA: Cohort x Arena half, F(1, 40) = 8.219, p<0.001; Unpaired two-sample t-test: Discrimination score, t = -3.256 p = 0.003)**. d.** Cre- cohorts prefer to interact with CS- whereas Cre+ mice spend comparable times interacting with CS+ and CS- (two-way ANOVA: Cohort x Stimulus mouse, F(1,40) = 7.429, p<0.01; Unpaired two-sample t-test: Discrimination score, t = 2.791, p = 0.01). Bonferroni post-hoc tests: * p<0.05, ** p<0.01, *** p<0.001. N = 5 female plus N = 5 male mice per cohort. Avpr1b-Cre+ and Amigo2- Cre+ numbers approximately balanced across cohort and sex with no significant effect of Avpr1b-Cre+ versus Amigo2-Cre+ genotype and sex observed.

**Extended Data Figure 5:**
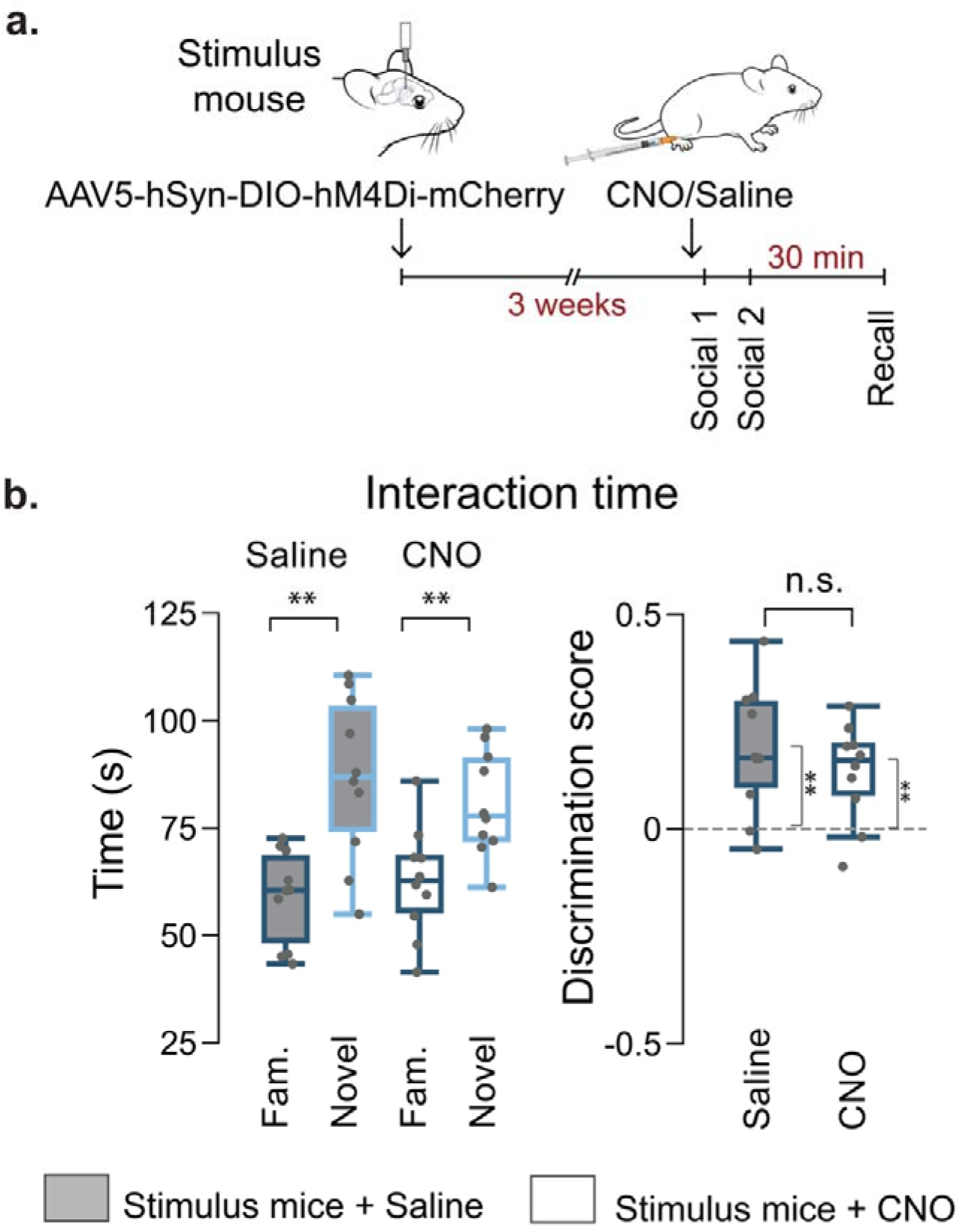
Social Memory assay with CA2-silenced stimulus mice. **a.** Social Memory assay with CA2-silenced stimulus mice. Male Avpr1b-Cre+ stimulus mice were injected in CA2 with Cre- dependent AAV2/5 hSyn.DIO.hM4D(Gi)-mCherry. After 3-4 weeks, CNO was administered to the stimulus mice 30 min prior to the first social memory encoding session (‘Social 1’). A second encoding session (‘Social 2’) follows immediately, with the position of the stimulus mice swapped. 30 mins later, a stimulus mouse, chosen at random, is replaced by a novel mouse (‘Recall’). N=5 subject mice. One week later (week 2), the procedure is repeated with saline injection administered to the same stimulus mice and a new cohort of subject mice (N=5). The same paradigm (CNO then saline injections in successive weeks) is then repeated with a different pair of stimulus mice and two cohorts of subject mice, resulting in a total of N = 20 male subject mice. **b.** Left: Time spent interacting with novel compared to familiar stimulus mice is comparable for CNO- and saline-injected stimulus mice cohorts. Two-way ANOVA: Injection type x Stimulus mouse, F(1,36) = 1.14, p>0.05. Bonferroni post-hoc tests: ** p<0.01. Right: Discrimination scores (novel versus familiar stimulus mouse) are comparable when stimulus mice have been administered CNO or saline (unpaired two-sample t-test: p = 0.39, t = -0.879). One-sample t-tests comparing discrimination score against zero: Stimulus mouse + saline, p = 0.0057, t = 3.603; Stimulus mouse + CNO, p = 0.0037, t = 3.882. Male Avpr1b-Cre stimulus mice and C57Bl/6J subject mice.

**Extended Data Figure 6:**
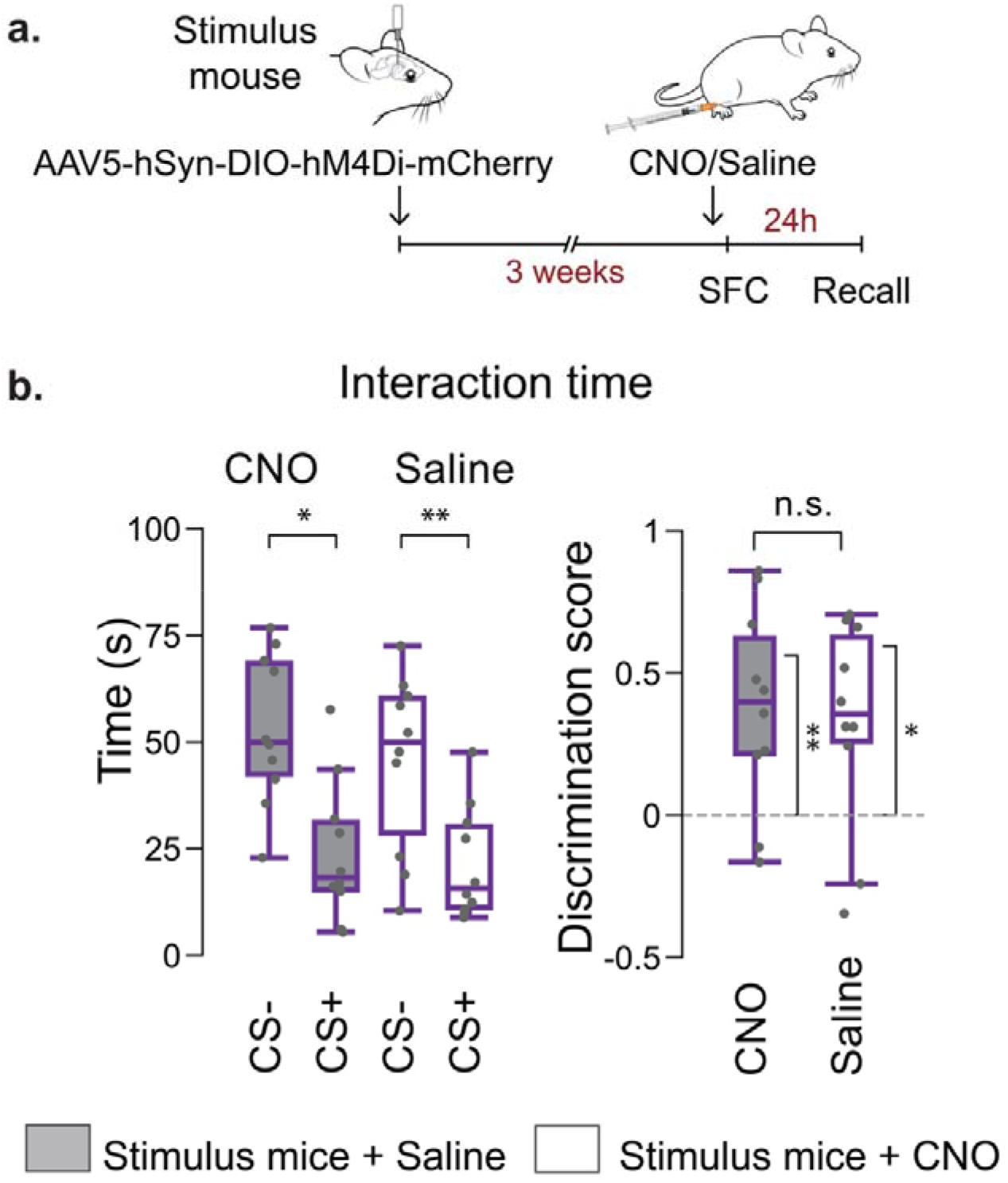
SFC assay with CA2-silenced CS+ and CS- stimulus mice. **a.** SFC assay with CA2-silenced stimulus mice. Male Avpr1b-Cre stimulus mice are injected in CA2 with Cre-dependent AAV2/5 hSyn.DIO.hM4D(Gi)-mCherry. After 3-4 weeks, CS+ and CS- stimulus mice receive CNO 30 min before SFC; N = 5 subject mice. The same stimulus mice are used for SFC one week later with saline injection and a new cohort of N= 5 subject mice. Then, the same procedure (CNO and saline) is repeated with a different pair of stimulus mice and two different cohorts of subject mice, resulting in a total of N = 20 male subject mice. **b.** Left: Subject mice spend a comparable time interacting with CS+ versus CS- stimulus mice during the recall session for CNO- and saline-injected stimulus mice. Two-way ANOVA: Injection type x Stimulus mouse, F(1,36) = 0.98, p>0.05. Bonferroni post-hoc tests: *p<0.05, ** p<0.01. Right: Discrimination scores (CS- versus CS+) are comparable when stimulus mice have been administered CNO or saline (unpaired two-sample t-test: p = 0.736, t = 0.342). One sample t-tests of discrimination score against zero: Stimulus mouse + saline, p = 0.02, t = 2.806; Stimulus mouse + CNO, p = 0.007, t = 3.403. Male Avpr1b-Cre stimulus mice and C57BL/6J subject mice.

**Extended Data Figure 7:**
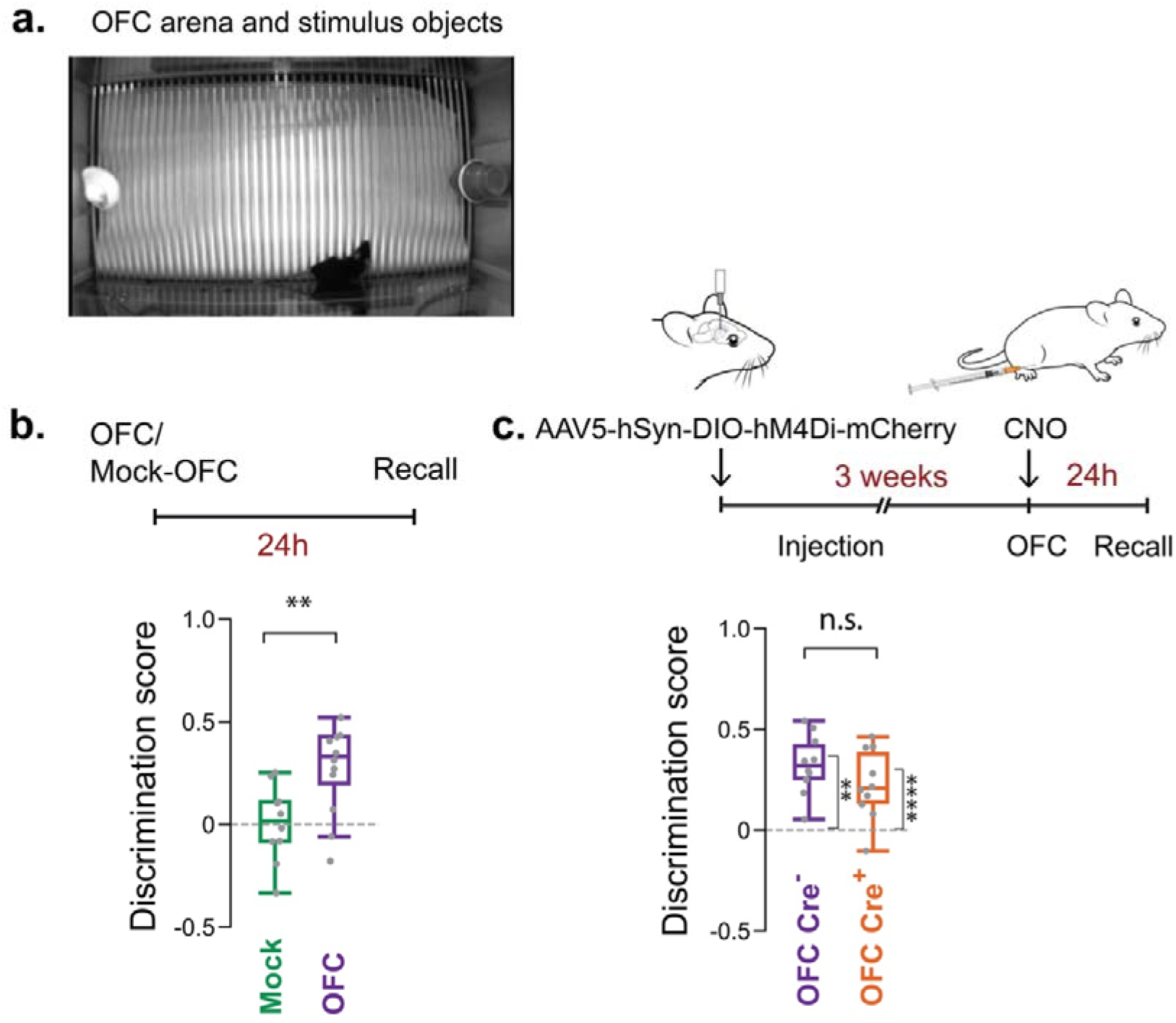
CA2 silencing does not affect Object Fear Conditioning. **a.** Object Fear Conditioning (OFC) assay. After 5 min habituation to the empty fear conditioning arena, mice receive a mild foot shock when interacting with a plastic duck (left) or two stacked red plastic cups (right), chosen at random as the CS+ object. 24 hours later, mice are re-exposed to the same objects in a novel recall arena. Mock-OFC cohorts undergo the same paradigm without footshock. **b.** OFC cohorts display a significantly greater preference for the safety- versus the threat-associated object than the Mock-OFC cohorts. Unpaired two-sample t-test: t = -3.048, p =0.006. N = 5 female mice plus 7 male mice, approximate sex-balanced across cohorts. **c.** Cre+ mice and Cre- littermate controls were injected in CA2 with Cre-dependent AAV2/5 hSyn.DIO.hM4D(Gi)-mCherry. After 3-4 weeks, mice underwent OFC 30 min after intraperitoneal CNO injection. Unpaired t-test: t = -1.380, p =0.184. One sample t-test against zero: Cre- cohort, t=6.94, p<0.0001; Cre+ cohort: t =4.11, p=0.0026. Avpr1b- Cre+ and Amigo2-Cre+ numbers balanced across cohort and sex. Avpr1b-Cre+ and Amigo2-Cre+ numbers balanced across cohort and sex with no significant effect of Avpr1b-Cre+ versus Amigo2-Cre+ genotype and sex observed.

**Extended Data Figure 8:**
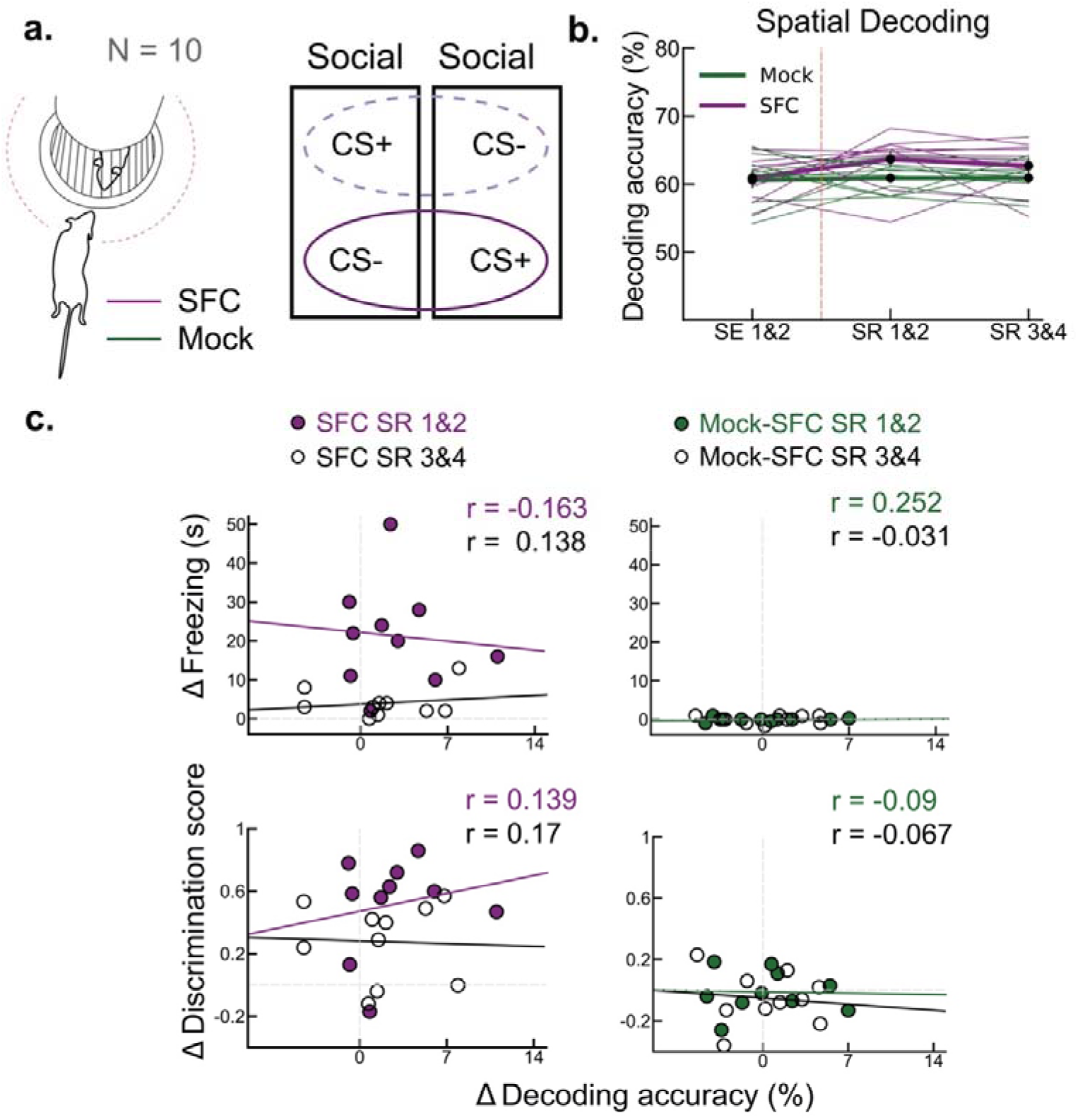
Linear decoding of spatial variables from CA2 population activity. **a.** Linear decoding analysis of position based on CA2 activity from calcium imaging during social/spatial interactions with the CS+ or the CS-, pooled across stimulus mice. Data around each cup balanced for mouse occupant by grouping data from two social trials. **b.** Spatial decoding accuracy is similar before (SE trial) and after (SR trials) SFC (purple). SFC and Mock-SFC (green) cohorts show similar spatial decoding accuracy throughout the paradigm, suggesting that the past threat-associated experience associated with a conspecific does not affect the representation of spatial location within the recall arena. Two-way ANOVA: Cohort x Session, F(2,54) = 1.284, p>0.05. **c.** Changes of spatial decoding are not correlated with change in freezing behavior (top graphs) or behavioral discrimination of CS+ from CS- (bottom graphs) for SFC (left graphs) or Mock-SFC (right graphs) cohorts as quantified by Spearman’s rank correlation coefficient, all correlations with p>0.05. N = 10 male mice with a total of 514 cells. N = 5 Avpr1b-Cre+ and 5 Amigo2-Cre+ with no significant effect of Avpr1b-Cre+ versus Amigo2-Cre+ genotype observed.

